# Mutation choice biases the structure of empirical fitness landscapes

**DOI:** 10.64898/2026.09.28.754994

**Authors:** Caelan Brooks, Michael M. Desai, Maryn O. Carlson

**Affiliations:** Department of Physics, Harvard University, Cambridge, MA, USA; Department of Organismic and Evolutionary Biology, Harvard University, Cambridge, MA, USA; NSF-Simons National Institute for Theory and Mathematics in Biology, Chicago, IL, USA; Department of Biology, University of Rochester, Rochester, NY, USA

## Abstract

Evolution can be thought of as taking place in a fitness landscape which maps each genotype to a fitness value. This abstraction has motivated efforts to quantify the statistics of random fitness landscapes with different amounts and types of interactions among loci, or epistasis. High-throughput experiments measuring the phenotypes of thousands of genotypes promise to bridge this theory with data. Yet thousands is still small relative to genotype space. Moreover, experiments often assay a highly non-random subset of genotypes. For example, a common approach is to assay all 2^*M*^ intermediates between ancestral and evolved sequences differing by *M* ≪ *L* mutations. Here, we ask how well the statistics of a landscape constructed under distinct “ascertainment” protocols recapitulate those of a random subset of the global landscape in several random landscape models. We show that, in most landscapes, selecting mutations that occurred along an adaptive trajectory inflates the variance explained by lower-order terms, reduces the number of local maxima, and increases the number of accessible paths relative to a random fitness landscape. Selecting the most beneficial or deleterious mutations instead distorts these statistics in ways that depend more critically on the amount of epistasis in the global landscape. We then use three large-scale mutagenesis datasets to “simulate” ascertainment in more realistic settings. In two of these, differences among schemes are likely explained by measurement noise and global nonlinearities; in the third, these differences at least partly reflect biased sampling of genuine interactions among loci.

## 1 Introduction

A fitness landscape maps all possible genotypes to fitness [Wright, 1932]. In theory, many features of evolution, such as its repeatability, can be predicted from the characteristics of a fitness landscape. Motivated by this abstraction, extensive experimental work has aimed to empirically characterize the structure of these high-dimensional landscapes in a diverse set of systems (see e.g. De Visser and Krug [2014] and Bank [2022] for recent reviews). However, generalizing across disparate experiments and systems poses several fundamental challenges.

One critical limitation is that even the largest empirical studies can assay only a minuscule fraction of genotype space. The extent to which any such local information about the fitness landscape reflects features of the global landscape is often unclear. For example, using a simulation-based approach, Draghi and Plotkin [2013] demonstrated that the magnitude and forms of epistasis among substitutions may vary along an adaptive walk, with epistasis becoming more prevalent as the walk proceeds. Agarwala and Fisher [2019] later showed that the distribution of epistatic effects for a broad class of fitness landscape models differs for genotypes obtained via uphill adaptive walks versus those simply selected based on their fitness differences from a shared ancestor. These findings suggest that the genetic backgrounds and mutations probed in experiments may often reflect atypical regions of a fitness landscape. More broadly, experimental design choices, including the environment, phenotype(s) assayed, fitness of the ancestral sequence [Reddy and Desai, 2021], and the number and choice of mutations [Szendro et al., 2013, Draghi and Plotkin, 2013, Bank et al., 2016, Pokusaeva et al., 2019], are likely to have a bearing on statistical features of empirical fitness landscapes.

Here, we study how the choice of which sites to mutate (referred to as ascertainment) distorts several frequently used summaries of fitness landscapes. We focus on a common type of combinatorial mutagenesis experiment (CME), in which the variant library consists of all 2^*M*^ possible intermediates between two sequences differing by *M* mutations [Fig. 1; Szendro et al., 2013]. In these experiments, we usually have *M*∼10, which is much smaller than the typical size of a protein (let alone a genome). Many of these experiments explore interactions among mutations presumed to have occurred along an adaptive trajectory [Weinreich et al., 2006, Khan et al., 2011, Moulana et al., 2022, Phillips et al., 2021, 2023]. For example, Khan et al. [2011] measured the fitness of all 2^5^ combinations of the first five mutations to fix in a long-term evolution experiment of *Escherichia coli* and Moulana et al. [2022] assayed landscapes involving all 2^15^ combinations of 15 mutations separating Omicron and Wuhan strains of the receptor binding domain of the SARS-CoV-2 spike protein. Other experiments have combined mutations with known individual functional effects [De Visser et al., 1997, Bank et al., 2016] or with known or hypothesized epistatic interactions [Wu et al., 2016, Salinas and Ranganathan, 2018, Johnston et al., 2024]. Still others combine alleles observed across orthologous sequences [Pokusaeva et al., 2019] or at high frequencies in natural populations [Schulz et al., 2025]. While previous studies have appreciated the potential effects of mutation choice on the statistics of empirical landscapes [Szendro et al., 2013, Bank et al., 2016, Pokusaeva et al., 2019, Agarwala and Fisher, 2019, Johnston et al., 2024], we still lack a comprehensive theoretical treatment of this topic.

**Figure 1.**
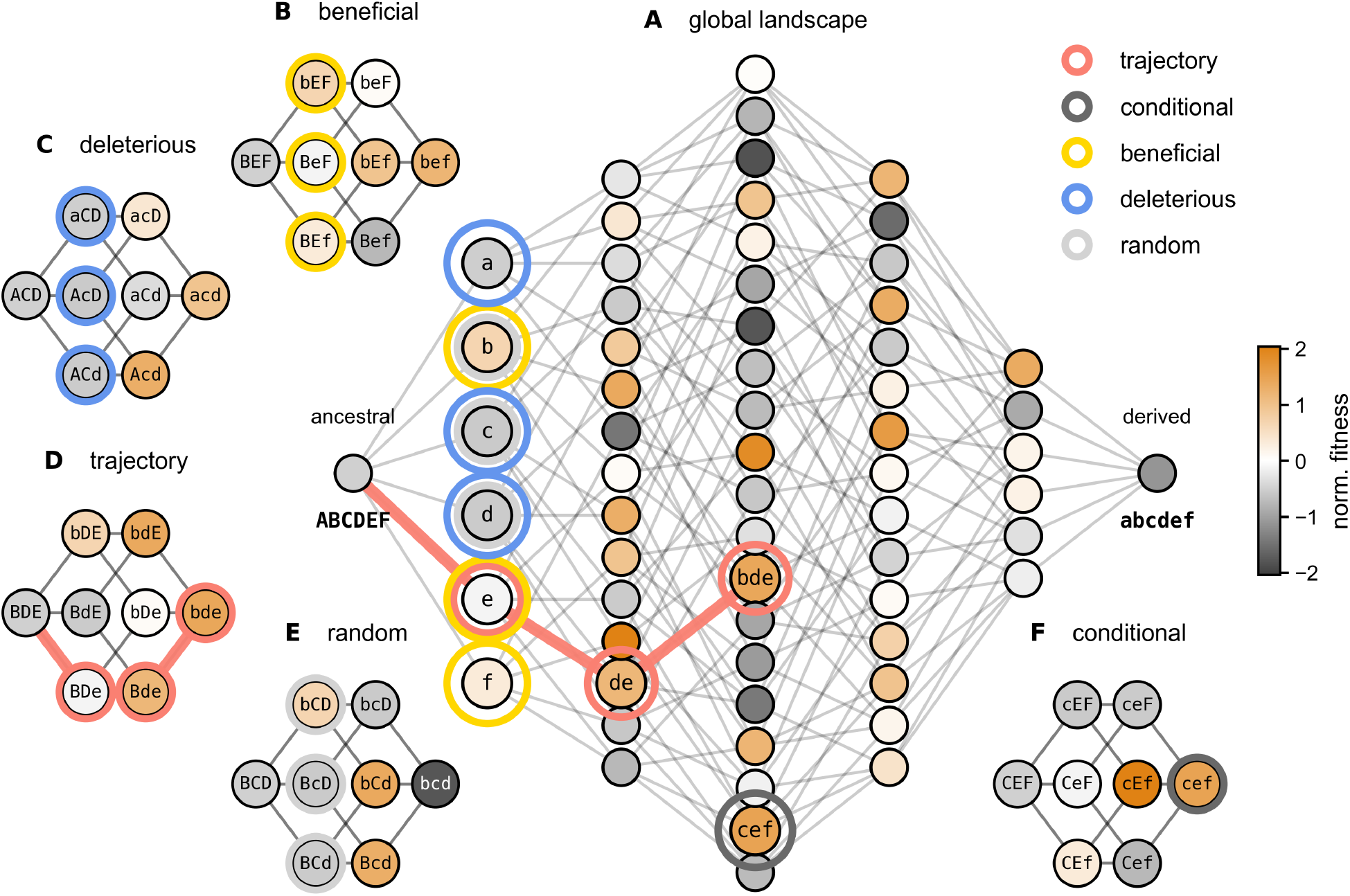
Schematic showing ascertainment schemes. **(A)** A global House of Cards (HoC) landscape (*L* = 6), with node color denoting fitness. The ancestral sequence is arbitrarily defined as ABCDEF, with mutant alleles indicated by lower case letters, a-f. Each single mutant is labeled with the corresponding mutated allele. **(B-F)** The global landscape was sub-sampled to generate five combinatorial libraries of size 2^*M*^ (*M* = 3) according to five ascertainment schemes: **(B)** beneficial; **(C)** deleterious; **(D)** adaptive trajectory; **(E)** random; and **(F)** conditional. The mutants which define each library are circled in **A-F.** The adaptive trajectory is indicated by bold salmon edges in **A** and **D**.

In this study, we aim to systematically analyze how ascertainment affects the statistics of CMEs. We begin by studying the effects of three types of ascertainment approaches in random landscape models, illustrated in Fig. 1B-D. Specifically, we analyze the statistics of CMEs in models where the mutations involved are chosen to be (1) the most beneficial mutations in a given ancestral sequence (beneficial), (2) the most deleterious mutations (deleterious), and (3) the mutations which occurred along an uphill, adaptive trajectory in the global landscape (trajectory). We compare the statistics of these “non-random” ascertained CMEs with two “random” ascertainment schemes (Fig. 1E-F): *M* uniformly sampled mutations (random), and *M* mutations such that the sequence at a distance of *M* mutations from the ancestor has the same fitness as that reached by an adaptive walk starting from the same ancestral sequence (conditional). We focus on the effects of ascertainment on three commonly employed statistics: the amount of variance explained by a regression model of order *n*, 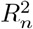, for *n* = 1, …, *M* ; the number of local maxima, *W* ; and the number of accessible paths, *P* [Szendro et al., 2013].

We first quantify the effects of ascertainment on these statistics in the House of Cards (HoC) landscape, an idealized model in which fitness differences are uncorrelated with genetic distance [Kingman, 1978]. The simple structure of the HoC landscape enables us to derive analytic approximations to the statistics of interest under each ascertainment scheme. We then consider two more realistic models which involve a tunable degree of distance-dependent correlations in fitness [Agarwala and Fisher, 2019]: the Rough Mount Fuji [RMF; Neidhart et al., 2014] and the LK [or NK; Kauffman and Weinberger, 1989] models.

We find that across the three classes of landscapes studied, non-random ascertainment of mutations often results in atypical landscape statistics with respect to a randomly ascertained CME. For example, in landscapes with sufficient amounts of epistasis, trajectory ascertainment inflates first-order effects and lower-order epistasis compared to random, resulting in higher 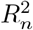 for all *n < M* . Beneficial and deleterious ascertainment yield more complex deviations from random, with over- or under-estimation of epistatic effects at a given order dependent on the structure of the global landscape. Ascertainment also leads to marked and epistasis-dependent differences in *W* and *P*, even when the 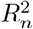 statistics are very similar across scenarios. These distortions, in part, follow trivially from the specifics of the ascertainment schemes. However, differences in *W* and *P* between conditional and trajectory ascertainment emphasize more subtle ways that the latter alters the curvature of a finite landscape.

Finally, we mimic the effects of ascertainment in empirical fitness landscapes by sub-sampling combinatorially complete experimental landscapes in which all possible mutations at a set of *M* sites have been measured. In two of these datasets, differences in the statistics among ascertainment schemes are likely described entirely by heteroskedasticity in measurement noise and the presence of global nonlinearities. However, in a third dataset, we find that differences in ascertainment schemes are, at least in part, more likely to reflect biased sampling of genuine epistatic interactions among loci.

## 2 Model

We first define global and local fitness landscapes using the Fourier decomposition formalism [Neher and Shraiman, 2011, Poelwijk et al., 2016, Agarwala and Fisher, 2019, Reddy and Desai, 2021]. We then introduce several statistics commonly used to summarize fitness landscapes [e.g., Szendro et al., 2013], and finally describe the five ascertainment schemes under consideration. Throughout, we focus on the regime in which the total size of the landscape is large compared to the size of the combinatorial library considered, as is typically the case in practice.

### 2.1 Global and local fitness landscapes

We model a genotype as a binary vector of length *L*, **x** ∈ {− 1, 1} ^*L*^, and let *F* (**x**) ∈ ℝ denote the fitness of **x**. The global fitness landscape is a mapping from the genotype space of 2^*L*^ sequences to ℝ. This landscape can be decomposed into contributions from interactions at each order:

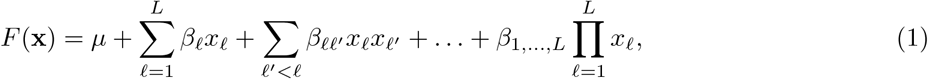

with interactions up to order *L*. Eq. 1 is the Fourier decomposition of the global fitness landscape, with coefficients given by weighted averages over all 2^*L*^ genotypes. Specifically, *µ* is the sample mean, and

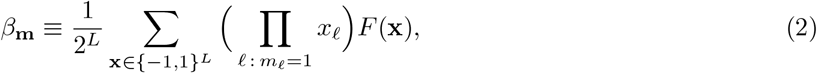

where **m** ∈ {0, 1} ^*L*^ is a binary vector with *m*≤*L* non-zero entries indicating the loci involved in the interaction [Weinberger, 1991, Hordijk and Stadler, 1998, Neher and Shraiman, 2011, Neidhart et al., 2013, Agarwala and Fisher, 2019]. Under complete sampling of the genotype space, the global variance in fitness is given by the sum of squares of coefficients at each order:

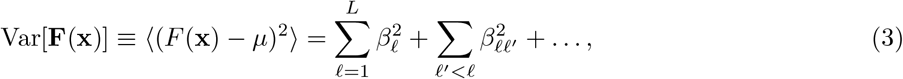

an important property of the Fourier decomposition that we will subsequently exploit to compute the variance explained by models with different orders of interactions.

Genotype space is often prohibitively large, even for modest *L*, so empirical studies can only explore a small subset of this space. A common experimental design is to generate a library of size 2^*M*^ consisting of all possible intermediates between two sequences, **x**_0_ and **x**_*M*_, that differ by *M* mutations. We refer to this type of experiment as a combinatorial mutagenesis experiment (CME) and the landscape it studies as a local fitness landscape within the larger global landscape. As with the global landscape, the local landscape studied in a CME can be described, without loss of information, by a Fourier decomposition with interactions up to order *M* . Under complete sampling of the size 2^*M*^ genotype subspace, the Fourier orthogonality relations and Eq. 3 hold (with summations over the *M* loci in the CME).

### 2.2 Local landscape statistics

Interpretation of high-dimensional fitness landscapes often relies on low-dimensional summary statistics. We use the following three statistics to quantify the effects of biased mutation ascertainment.

#### 2.2.1 Variance explained

For a given combinatorial library, it is common practice to ask to what extent a regression model of a given order explains the observed phenotypic variance [Szendro et al., 2013, Phillips et al., 2023, Park et al., 2024]. For example, for order *n* = 2, one fits the following regression model: 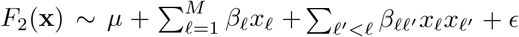,where *µ* is the sample mean and *ϵ* is a normally distributed error term. The variance 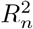, explained by a model of order *n*, 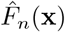, is then defined as

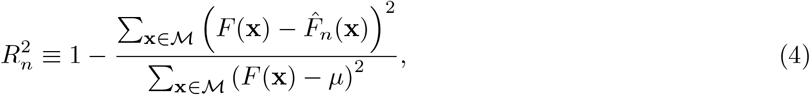

where ℳ is the set of 2^*M*^ genotypes in the CME and 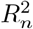 is the predicted fitness of sequence **x** derived from fitting a model with up to order *n* (*n*≤ *M* ) interactions.

In the absence of measurement noise—an assumption which allows us to isolate the effects of ascertainment on the statistics of empirical landscapes—and when all 2^*M*^ genotypes are sampled, 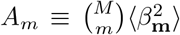 is directly related to the amplitude spectrum, *A*_*m*_ (*m* = 1, …, *M* ), of the local fitness landscape [Szendro et al., 2013]. Each element of *A*_*m*_ is defined as the sum of squared coefficients of order *m*, 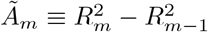, where, as above, **m** ∈ {0, 1}^*M*^ with |**m**| = *m*. For example, for *m* = 2,

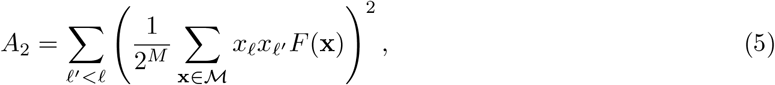

and so on for *m* ∈ ℕ+, *m* ≤ *M* . The orthogonality of the Fourier decomposition (see Eq. 3) implies

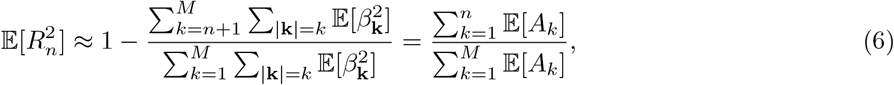

where the expectation of the ratio in Eq. 6 is approximated as a ratio of expectations. Note that the expectation in Eq. 6 is over realizations of the fitness landscape. We use the bracket notation, ⟨·,·⟩ to denote an average over the coefficients, and thus fitness values, within a single realization of the landscape. We are also sometimes interested in the fraction of phenotypic variance explained by interactions of a given order, denoted by 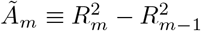 for *m* = 1, …, *M*, with 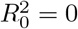.

#### 2.2.2 Number of local maxima

The number of local maxima, *W*, is often used as a measure of the “ruggedness” of a fitness landscape. *W* is computed for a given landscape by considering the single mutation neighborhood of each member of a combinatorial library, 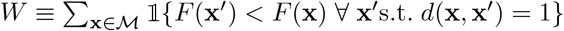, where *d*(·, ·) denotes the Hamming distance.

#### 2.2.3 Pathway accessibility

The number of accessible uphill paths, *P*, is the number of paths from the ancestral to the fully derived sequence in which each step increases fitness. Formally, we have 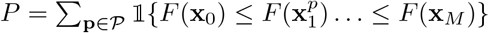 where **p** denotes a given path of *M* + 1 sequences, 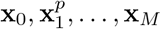, in the set *P* of *M* ! possible paths.

### 2.3 Biased sub-sampling of genotype space

CME libraries are often constructed from sets of highly atypical mutations. We consider three ways of ascertaining mutations inspired by common experimental strategies (Fig. 1). In the (1) beneficial and (2) deleterious scenarios, the library is constructed from the *M* most beneficial and *M* most deleterious mutations in the ancestor, respectively. Under (3) trajectory ascertainment, the *M* mutations are those which occur along an uphill adaptive trajectory. We assume that at each step in the adaptive trajectory the subsequent mutant is selected uniformly from among the available beneficial mutations in the global landscape.

We compare the local fitness landscapes created in these three scenarios to those constructed from libraries involving (4) random mutations (uniformly sampled from all *L* possible loci) and (5) conditional libraries for which the ancestral and fully derived sequence have the same fitness as in a matched trajectory scenario. The random scenario exclusively captures the effects of downsampling from *L* to *M* loci. In the trajectory scenario, the fitness of the derived sequence is often much greater than that of the ancestral sequence (*F*_*M*_ ≫*F*_0_), trivially enriching for accessible paths relative to a random library. Thus, the conditional scenario isolates the more subtle effects of trajectory ascertainment on the statistics of a CME.

We assume throughout that the ancestral background — shared across ascertainment scenarios — is sampled uniformly from all 2^*L*^ possible genotypes in the global landscape. In practice, the ancestral background may also be atypical, a subject we return to in Section 6.

## 3 House of Cards fitness landscapes

We first study the effects of ascertainment on CMEs when the global fitness landscape is described by the House of Cards (HoC) model [Kingman, 1978]. The HoC landscape is defined as one in which the fitness of each genotype, *F* (**x**), is an independent and identically distributed (*iid* ) sample from a normal distribution. That is, 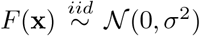. Without loss of generality, we let *σ*^2^ = 1. The HoC landscape represents a high-epistasis limit of statistical landscape models in which there is no correlation between the fitness of neighboring genotypes.

In the following sections, we describe the effects of each ascertainment scheme on the three focal statistics, 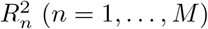, *W*, and *P* . We also compare these results to simulations, with both illustrated in Fig. 2.

**Figure 2.**
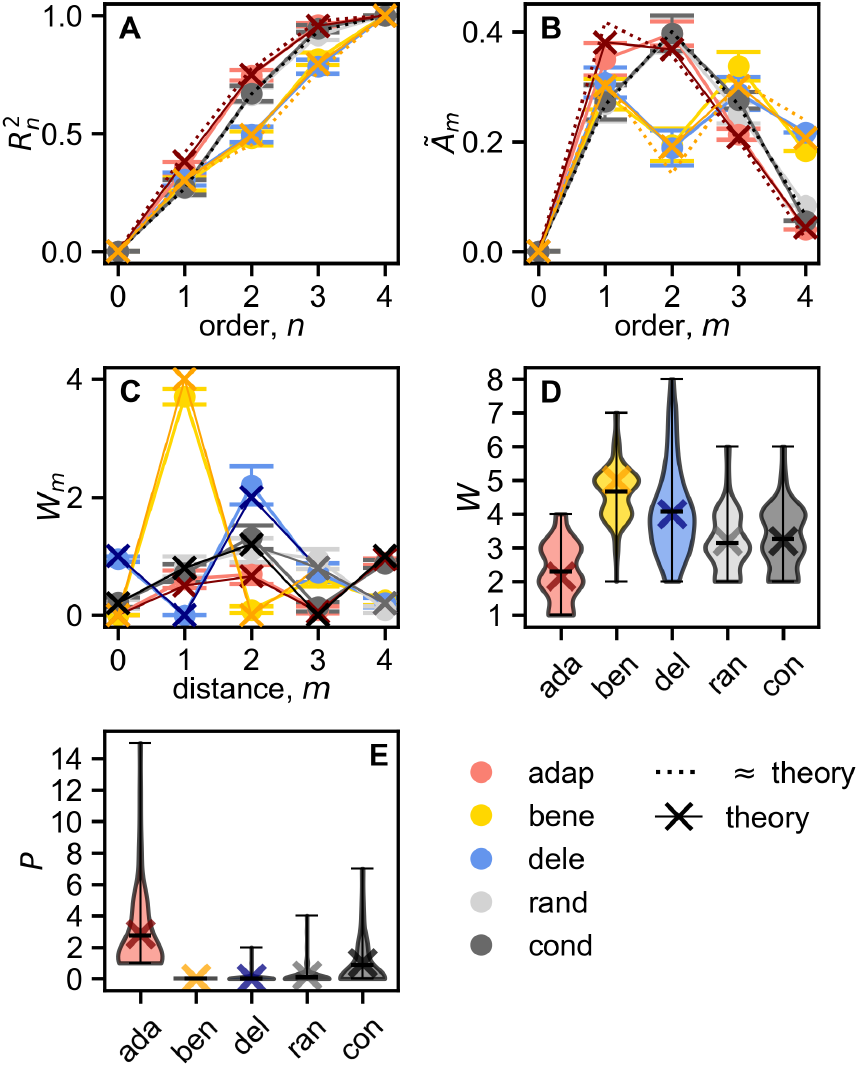
Statistics of ascertained libraries in House of Cards (HoC) landscapes. **(A)** Variance explained by models of order *n*,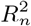, *n* = 0, …, *M*, for combinatorial libraries with *M* = 4 for the adaptive trajectory (salmon), beneficial (yellow), deleterious (blue), random (gray), and conditional (dark gray) ascertainment schemes in HoC landscapes with *L* = 100, averaged over *N* = 100 simulations. Error bars represent 95% confidence intervals. Theoretical expectations are shown with ×’s. **(B)** The normalized amplitude spectrum, 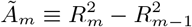, with all otherwise as in **A.** Dotted lines in **A**-**B** denote expectations for the statistics under the assumption that the atypical mutants in these libraries are *iid* samples from a distribution with mean *µg* = 4 and 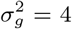, and deleterious single mutants from a distribution with mean −*µ*_*g*_ and variance 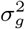. **(C)** The average number of local maxima as a function of distance, *m*, from the ancestral sequence, *W*_*m*_. **(D)** Violin plots of the number of local maxima, *W*, and **(E)** accessible paths, *P*, across ascertainment schemes. Black hashes in **D**-**E** denote the maximum, mean, and minimum, respectively.

### 3.1 Variance explained by interactions of a given order

We begin by exploiting the relationship between 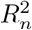 and *A*_*m*_ (Eq. 6) to solve for the former in each ascertainment scenario in CMEs consisting of *M* mutations.

Under random ascertainment in the HoC, *A*_*m*_ takes a simple form, with 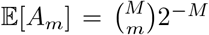, shown in Fig. 2B for *M* = 4 (see Fig. S1 for *M* = 2, 4, 6). *A*_*m*_ is fully determined by the number of combinations of *m* loci [Neher and Shraiman, 2011, Neidhart et al., 2013, Agarwala and Fisher, 2019]. This result follows from the fact that the fitness values of distinct sequences are *iid* (Section S1.1). Thus, under random ascertainment, 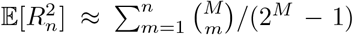, where the denominator is the expectation of the sample variance (Fig. 2A).

Conditional ascertainment, with the extreme fitness of the endpoint sequence, *F*_*M*_, inherited from an adaptive walk, violates the *iid* assumption. However, ignoring the correlation between the ancestral and endpoint fitness values, an *O*(2^*−*2*M*^ ) correction, conditional CME yields precisely the same *A*_*m*_ as a random CME (Fig. 2, Section S1.2).

The three non-random ascertainment schemes, however, result in systematic distortions of *A*_*m*_, and thus of 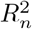 . Before more explicitly modeling the distributions of the atypical mutants in each non-random CME (see below and Section S1), we consider a simplified scenario in which the fitness values of the *M* beneficial and trajectory mutants are sampled *iid* from a distribution *G* with mean *µ*_*g*_ ≥ 0 and 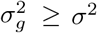. We likewise assume that the deleterious mutants are sampled from a distribution, 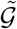, with mean *µ*_*g*_ and variance 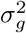. Thus, in each scenario, we are violating the second feature of random ascertainment, identical means, while retaining the first, independent fitness values. This simplification will allow us to obtain analytic, interpretable approximations for the 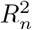 statistics, while qualitatively capturing the effects of choosing mutations with arbitrarily beneficial or deleterious effects.

In the beneficial case, we solve for *A*_1_ by finding the expected value of an order one squared coefficient,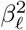,

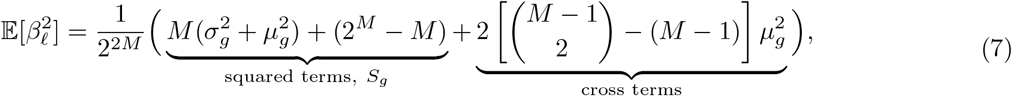

where the partitioning of the beneficial single mutants based on their genotype states at locus *l* results in a non-zero contribution from the cross terms (see Eqs. 2 and 5). The expected value of *A*_1_ is found by multiplying Eq. 7 by *M* . To generalize to arbitrary *m*, one need only consider how each interaction of order *m* partitions the beneficial mutants, yielding

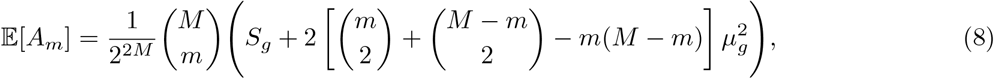

where *S*_*g*_ denotes the contribution from the squared terms to each element of *A*_*m*_ (Eq. 7; Fig. 2B). (We employ the convention that 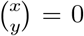 for *x < y*.) Expected 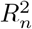 follows by summing the elements of *A*_*m*_ for *m* = 1, …, *n* and normalizing by the expected sample variance, 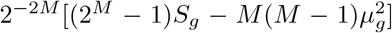. For *µ*_*g*_ = 0, regardless of 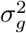, we recover the random HoC result. In Section S1.5, we relate Eq. 8 to a microscopic specification of the HoC model.

It is easily seen from Eq. 8 that deleterious ascertainment will yield the same 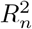 statistics as beneficial. This result does not, however, rely on our simplified distributional assumptions, only the symmetry imposed by sampling the ancestral sequence uniformly from the global landscape (Fig. 2A-B). In practice, the distributions of deleterious and beneficial mutants likely diverge in a manner that depends on the fitness of the ancestral sequence (see Section 6).

In the trajectory case, as in the beneficial scenario, the partitioning of atypical mutants determines the perturbed amplitude spectrum. However, the fact that the adaptive mutants are at distances of *d* = 1, …, *M* from the ancestral sequence results in a distinct partitioning and thus spectrum. To see this, consider a single locus *l* and assume that the mutations are ordered from 1 to *M* in accordance with their occurrence in the trajectory. The trajectory mutants with *i* ≥ *l* will share a genotypic state at locus *l*, partitioning them into two sets: the *M* − *l* + 1 with the mutation and the *l* − 1 without the mutation. An expression for the expected value of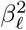 and thus *A*_1_ immediately follows:

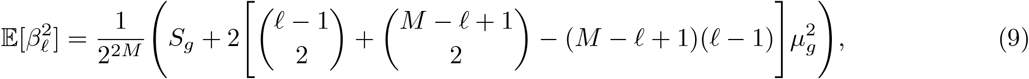

where *S*_*g*_ was defined in Eq. 7, and the rightmost terms (multiplying 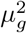) represent the cross terms. By averaging over *l*, we find

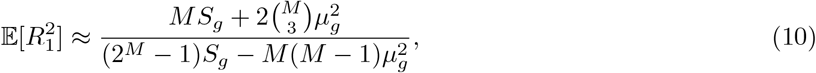

where, under our simplifying assumptions, the denominator is equal to that under beneficial choice (Fig. 2A-B).

At higher orders, the partitioning is more complex. In Section S1, we derive an implicit expression for *A*_*m*_ by enumerating the partitions at each order (dotted lines, Fig. 2A-B). For *M* = 4, inflation of the 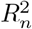 statistics can be attributed primarily to increased 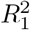 (Eq. 10). This is in contrast to the beneficial scenarios, where *m* order and *M* −*m* order interactions equivalently partition the atypical mutants. This symmetry results in inflation of both lower and higher order epistasis for beneficial relative to random ascertainment, with depression of epistasis at intermediate orders.

We achieve more accurate expectations for 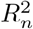 in the beneficial scenario by modeling the fitness of the single mutants as the order statistics of *L iid* standard normal random variables [Blom, 1958, David and Nagaraja, 2003]. In the trajectory scenario, we use estimates of the moments of the mutants along an uphill adaptive trajectory to refine expressions for 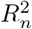 (solid lines, Fig. 2A-B; Section S1).

As *M* increases, deviations from random in all non-random scenarios diminish as the atypical mutants comprise a decreasing fraction of the assayed mutants (*M*· 2^*−M*^ →0) and the fitness values of mutants in the library are otherwise uncorrelated.

### 3.2 Local maxima

We next consider how the expected number of local maxima, *W*, depends on ascertainment. Under random ascertainment, we have E[*W* ^rand^] = 2^*M*^ */*(*M* + 1) as each sequence is equally likely to be the maximum among its *M* neighbors [Kauffman and Levin, 1987]. The number of local maxima at a given distance from the ancestor, *W*_*m*_, is thus proportional to the number of locus combinations at each order, 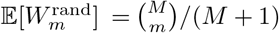, akin to *A*_*m*_ (Fig. 2C-D).

As with 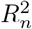, conditional ascertainment yields approximately the same *W* as that of random (Fig. 2D). For modest *M*, we can assume that an adaptive trajectory achieves sufficiently high fitness, *F*_*M*_, such that the probability *F*_*M*_ is a local maximum is approximately one, i.e., Φ[*F*_*M*_ ]^*M*^≈ 1, where Φ[· ] is the cumulative distribution function (CDF) of a standard normal random variable. Yet, at distances greater than one mutation from **x**_*M*_, the probability that a given mutant is a local maximum is the same as under random ascertainment. We therefore have, 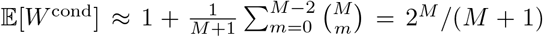, identical to that of random. Thus, while the conditional scenario alters expected *W*_*m*_ for *m* = *M, M*− 1, E[*W* ^cond^] = E[*W* ^rand^] (Fig. 2C).

Non-random ascertainment, however, has a more profound impact on both *W* and *W*_*m*_ in a CME: As trajectory mutants are ascertained under the assumption that *F*_0_ ≤*F*_1_ ≤… ≤ *F*_*M*_, only *F*_*M*_ among the trajectory mutants has a non-zero probability of being a local maximum. In addition, a non-trajectory mutant with one or more trajectory neighbors will be a local maximum with reduced probability relative to random, as long as E[*F*_*i*_] *>* 0. At a given genetic distance *d* from the ancestor 1 ≤ *d < M*, there are *M− d−*1 non-trajectory mutants with *F*_*d−*1_ as a neighbor and *d−* 1 non-trajectory mutants with *F*_*d*+1_ as a neighbor. For 1≤ *d < M*, there is one non-trajectory mutant with two trajectory neighbors. We use these observations to approximate expected 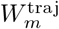 and *W* ^traj^ (Fig. 2C-D; Section S1.3). We observe that trajectory ascertainment results in a “smoother” landscape with fewer local maxima (Fig. 2D), consistent with inflated first-order effects and lower-order epistasis as indicated by 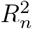 (Fig. 2A). Importantly, this smoothing effect results in deviations from conditional in terms of *W*_*m*_ (for 1 *m M* 2) and *W* (Fig. 2C-D).

Beneficial ascertainment has the opposite effect, inducing a more rugged local landscape (Fig. 2). Let *F*_(1)_, …, *F*_(*M*)_ denote the fitness values of the beneficial single mutants. Each of the single mutants is a local maximum with probability Φ[*F*_(*i*)_]^*M−*1^, as each mutant has *M* neighbors, one of which is the ancestral sequence, with *F*_(*i*)_ ≥ *F*_(0)_ by definition. When the number of loci *L* is large, Φ[*F*_(*i*)_] ≈ 1 for *M* ≪ *L*, and E[*W* ] ≈ *M* . Consequently, each of the 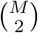 mutants at a distance of two mutations from **x**_0_ is unlikely to be a local maximum, and 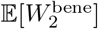. At a distance *d >* 2, the neighborhood of each mutant is identical to that of the random HoC, and thus the probability that a given mutant is a local maximum is (*M* + 1)^*−*1^ (Fig. 2C). Combining these results, we find

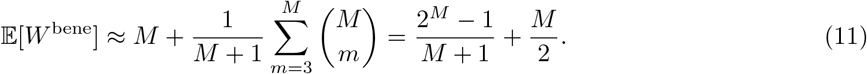

Therefore, for large *M, W* ^bene^ typically exceeds *W* ^rand^ by approximately *M/*2 local maxima (Fig. 2D).

For the deleterious case, we abuse notation and let *F*_(1)_, …, *F*_(*M*)_ denote the fitness values of the deleterious mutants. For *M*≪ *L*, 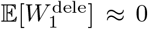. The extreme small values of the *F*_(*i*)_ in turn increase the probability that mutants at a distance of *d* = 2 are local maxima. For *d >* 2, the probability that a mutant is a local maximum is equal to that of random choice. Combining these considerations,

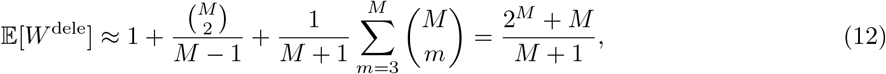

and thus, deleterious choice results in approximately one additional local maximum when compared to random (Fig. 2C-D). Despite identical 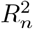 statistics to beneficial, deleterious ascertainment yields distinct *W*_*m*_ and *W*, with the latter approximating *W* ^rand^ for modest *M* .

### 3.3 Pathway accessibility

In a random local landscape, the probability that a given path *p* from **x**_0_ to **x**_*M*_ for *F*_*M*_ *> F*_0_ is accessible is given by 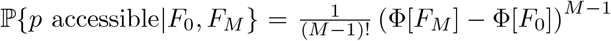, where *F*_*M*_ and *F*_0_ are the fitness values of the derived and ancestral sequences [Krug and Oros, 2024]. The expected value of *P* ^rand^ is given by integrating over both *F*_0_ and *F*_*M*_, yielding E[*P* ^rand^] = 1*/*(*M* + 1).

Conditional ascertainment trivially alters *P* by increasing the probability that a given path is accessible due to the fact that *F*_*M*_≫ *F*_0_. Integrating over *F*_0_ and assuming *F*_*M*_ ≫ 0, we find that E[*P* ^cond^] ≈ 1. Thus, unlike *P* ^rand^, which scales inversely with library size, *P* ^cond^ does not depend on *M* (Fig. 2E).

Trajectory ascertainment further increases *P* beyond simply conditioning on *F*_*M*_ ≫ 0 (Fig. 2E). By definition, the adaptive trajectory represents an accessible path. In addition, the inclusion of any subset of trajectory mutants in a given path will alter the likelihood that the path is accessible. To find the expected value of *P* ^traj^, we thus consider the number of paths which involve each possible subset of trajectory mutants (Section S1.3).

Beneficial and deleterious ascertainment, however, trivially reduce the expected number of uphill paths to approximately zero (Fig. 2E): Deleterious single mutants are by definition less fit than the ancestral sequence, precluding uphill paths for *M*≪ *L* and large *L*. Likewise, in this regime, combinations of beneficial mutations are unlikely to yield fitness values greater than those of the corresponding single mutants.

## 4 Alternative fitness landscapes

While the HoC model provides a tractable system for studying the effects of ascertainment, it represents a high-epistasis extreme case where there is no correlation in fitness between neighboring genotypes. Here, we explore the extent to which the conclusions derived from the HoC model hold in two more biologically realistic landscape models which involve some degree of distance-dependent divergence: the Rough Mount Fuji [RMF; Aita et al., 2000, Neidhart et al., 2014] and the LK [more commonly NK; Kauffman and Weinberger, 1989] landscapes.

In an RMF landscape, the fitness of a sequence **x** ∈ {−1, 1}^*L*^ is the sum of additive and HoC components:

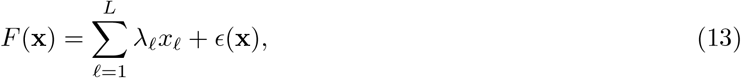

where 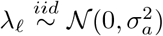 and 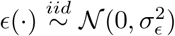 [Neidhart et al., 2014, Agarwala and Fisher, 2019]. The original formulation of the model is recovered by letting *λ*_*l*_ = 1 for all *l* [Aita et al., 2000]. *By tuning the relative magnitudes of σ*_*a*_ and *σ*_*ϵ*_, the RMF model interpolates between additive (*σ*_*a*_ *>* 0, *σ*_*ϵ*_ = 0) and HoC landscapes (*σ*_*a*_ = 0, *σ*_*ϵ*_ *>* 0). Depending on the relative contributions of the additive and HoC components to phenotypic variance, a mutation may be more likely to have a large effect due to its first order effect, 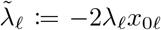, where *x*_0*l*_ is the state of the ancestral sequence at locus *l*, or its value of *ϵ*, i.e., its interactions with other loci. We thus study ascertainment in the RMF as a function of *σ*_*ϵ*_, with *σ*_*a*_ = 1.

In an LK landscape, fitness is instead determined by contributions from *L* locus neighborhoods of size *K* [Kauffman and Weinberger, 1989]. *For a given locus i*, each of the 2^*K*^ genotype combinations maps to a unique value, 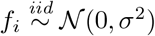,

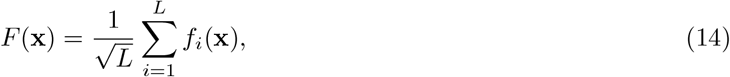

where implicitly, *f*_*i*_(·) is a function of **x**∩ℬ_*i*_, and ℬ_*i*_ is the size *K* neighborhood of locus *i*. Without loss of generality, we let *σ*^2^ = 1. As in Neidhart et al. [2013], we consider a simplified model in which locus *i* does not necessarily participate in its respective neighborhood, ℬ_*i*_.

Like the RMF, the LK model contains both the additive (*K* = 1) and HoC (*K* = *L*) landscapes [Kauffman and Levin, 1987, Kauffman and Weinberger, 1989]. For intermediate *K*, a mutation’s effect is determined by the number of neighborhoods in which it participates, *B*_*i*_, as well as the fitness contributions of those neighborhoods, *f*_*i*_. Interactions among loci are determined by the number of neighborhoods that they share. We denote the number of shared neighborhoods between two mutations *i* and *j* as *B*_*ij*_. We will use these three metrics to understand how ascertainment perturbs local landscape statistics in LK landscapes as a function of *K*, for fixed *L*.

### 4.1 Variance explained

Under random ascertainment in the RMF, the relative magnitudes of *σ*_*a*_ and *σ*_*ϵ*_ solely determine *A*_*m*_, and thus 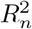 . In a CME with *M* mutations,

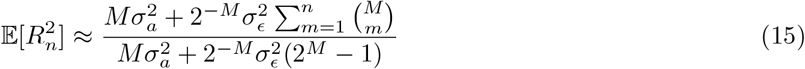

[Neidhart et al., 2013, Section S3.1]. Even under random ascertainment, the 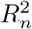 statistics exhibit a non-monotonic dependence on *M* (Fig. S3).

As in the HoC, in the global LK landscape, *A*_*m*_ is proportional to the number of interactions of a given order, 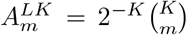 . However, here, 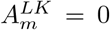 for all *m > K* as interaction order cannot exceed *K* [Neidhart et al., 2013]. Under random downsampling, 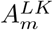 is less compact,

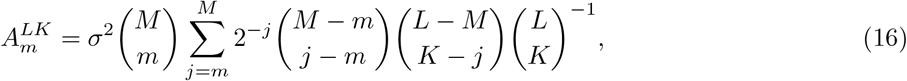

a result derived in Section S4.1. Briefly, all neighborhoods containing at least *m* of the *M* loci will contribute to an interaction of order *m*. The probability of, for example, *j* of the *M* loci falling in neighborhood *i* is given by the hypergeometric distribution — the rightmost three terms in the summand. The contribution of a neighborhood to *A*_*m*_ scales exponentially with the number of ascertained mutations in that neighborhood, *j*; neighborhoods with fewer (but≥ *m*) mutations contribute more to *A*_*m*_.

We see immediately that unlike in the RMF and HoC models, 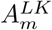 (and thus 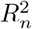 ) depends on the size of the global landscape, *L*. In addition, for a fixed *m*, 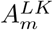 is potentially a non-monotonic function of *K*, whereas the simple structure of the RMF landscape guarantees that the normalized spectrum, 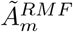, is an increasing function of *σ*_*ϵ*_ for *m*≥ 2 (Fig. S4).

Yet, ascertainment generally manifests similarly in both landscapes (Fig. 3). To ground our observations, we start with the additive limit where ascertainment has no effect on the *R*^2^ statistics: 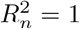 for all *n* ≥1. As one approaches the HoC limit by increasing *σ*_*ϵ*_ and *K*, 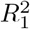 necessarily decreases while *Ã*_2_ increases (at least initially) across all schemes (Fig. 3A-D).

**Figure 3.**
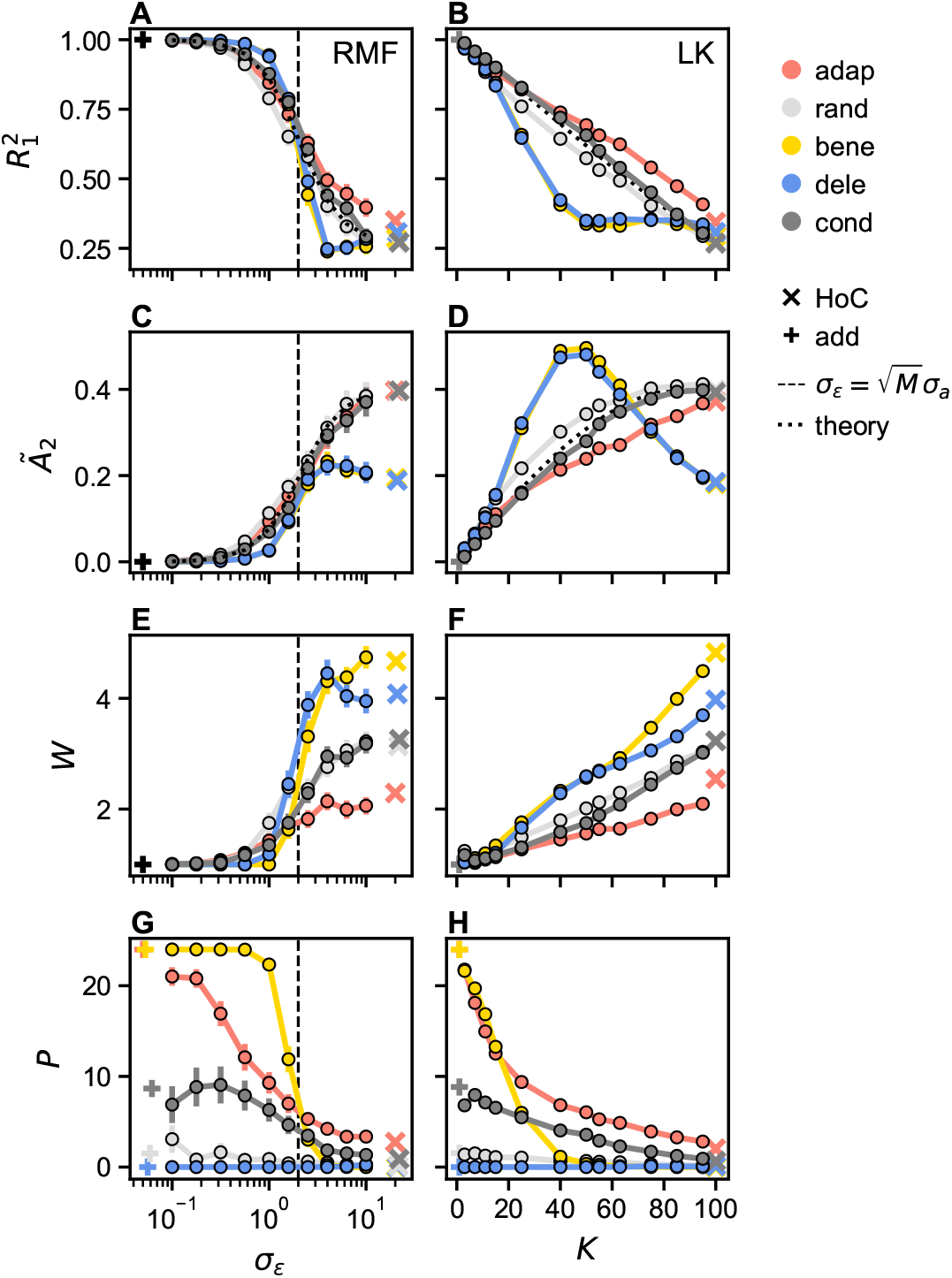
Effects of ascertainment on the statistics of tunable landscapes. Statistical summaries of ascertained fitness landscapes with *M* = 4 loci sub-sampled from *N*_rmf_ = 100 simulated Rough Mount Fuji (RMF; left column) and *N*_LK_ = 1,000 simulated LK (right column) landscapes with *L* = 100 loci. In the RMF landscapes, 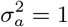 and *σ*_*ϵ*_ varies over two orders of magnitude (*x*-axis). In the LK landscapes, *K* varies from one to *L* = 100 (*x*-axis). Average variance explained by **(A-B)** first order, 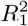, and by **(C-D)** second order, 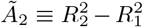, interactions, respectively, as functions of *σ*_*ϵ*_ and *K* for each ascertainment scenario (colored points). The dotted lines denote theoretical expectations for each quantity (Eqs. 15 and 16); the dashed vertical lines in the left column denote where 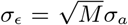. The same statistics estimated from simulations of a House of Cards (HoC) landscape are denoted by ×’s, while those computed under an additive landscape are denoted by +’s. The average **(E-F)** total number of local maxima, *W*, and **(G-H)** number of accessible paths, *P*, likewise as functions of *σ*_*ϵ*_ and *K*, with colors and symbols as in **A**-**B**.

In both models, trajectory ascertainment results in inflated *R*^2^ relative to random and conditional only as the amount of epistasis increases, with deviations more pronounced in the LK. For larger *K*, we observe that mutations which occur along an adaptive trajectory share fewer neighborhoods with one another than random (Δ*B*_*ij*_ *<* 0) and conditional mutations, in part, because they are less connected in the global landscape (Δ*B*_*i*_ *<* 0; Fig. 5A,C). Thus, in the LK, trajectory ascertainment enriches for non-interacting sets of mutations more effectively than in the RMF landscape, resulting in larger *R*^2^ deviations from random compared to the RMF.

In the RMF, at small *σ*_*ϵ*_, and if *F*_*M*_ values are large enough, a sufficient fitness increase in the conditional library can only be obtained by sampling mutations with similarly large additive effects (Fig. 4E). For intermediate *σ*_*ϵ*_, however, comparable fitness gains can be achieved with a large value of *ϵ* for the fully derived sequence and more modest additive effects. As the HoC component begins to dominate, 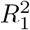 is inflated in trajectory relative to conditional due to the considerations described in Section 3.

**Figure 4.**
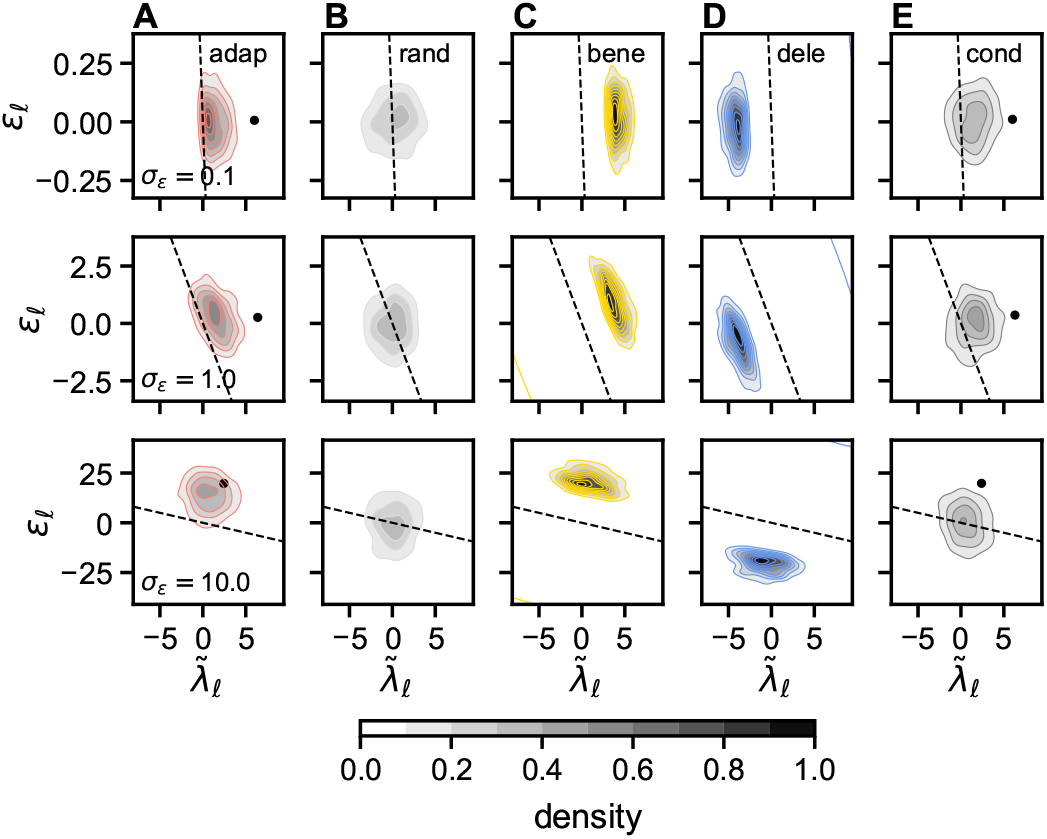
Effects of ascertainment on mutation features in Rough Mount Fuji landscapes. Combinatorial libraries of size 2^*M*^, with *M* = 4, were ascertained from Rough Mount Fuji (RMF) with varying *σ*_*ϵ*_, with corresponding summary statistics shown in Fig. 3. Contour plots **(A-E)** represent the HoC (*ϵ*_*l*_, *y*-axis) and additive (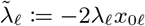, *x*-axis) components of each mutation for each ascertainment scheme with *σ*_*ϵ*_ = .1 (top), *σ*_*ϵ*_ = 1 (middle), and *σ*_*ϵ*_ = 10 (bottom), respectively. For the trajectory scenario (shown in **A**), the distributions correspond to the mutants which occurred along the adaptive trajectory. All other subplots show the HoC components of the single mutants with respect to the ancestral sequence (the additive component is a property of the mutation). In addition, in **A** and **E**, the mean fitness components of the fully derived sequences under trajectory and conditional ascertainment are denoted by black points, respectively. The dashed lines denote where 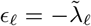.

In both landscapes, 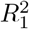 and *Ã*_2_ under beneficial and deleterious ascertainment exhibit a non-monotonic dependence on the extent of epistasis (Fig. 3A-D). Deleterious ascertainment in both landscapes yields precisely the same behavior as beneficial, so we refer only to beneficial henceforth. In the RMF, beneficial ascertainment first induces marked inflation of 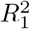 relative to random ascertainment, before exhibiting depressed 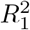, and finally again inflated 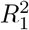 in the HoC limit. In contrast, in the LK, beneficial ascertainment does not initially exhibit inflated 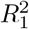 (Fig. 3B). This discrepancy is explained by differences between the beneficial mutations in each landscape. In the RMF, when 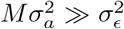, beneficial mutants are enriched for large additive effects (Fig. 4C). Whereas, in the LK, for 1≤*K*≪*L*, beneficial mutations tend to participate in (1) more neighborhoods in the global landscape and (2) to share more (large effect) neighborhoods with each other (Fig. 5). The latter, in particular, results in more epistasis among beneficial mutants relative to random loci. For the same reasons, second-order interactions explain relatively less variance in the beneficial RMF versus LK libraries for small *σ*_*ϵ*_ and *K*, respectively (Fig. 3C-D).

As 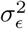 increases relative to 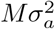, the most beneficial mutants in the RMF are instead associated with large values of *ϵ*, and, equivalently, more epistasis, with 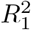 dipping below that of random (Fig. 4C; Section S3.2). Beneficial 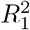 likewise decreases with increasing *K* until *K* ∼ *L* at which point it is relatively constant and approximately equal to its HoC limit. In contrast to the RMF model, 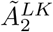 exceeds that of random at intermediate *K*, where the difference between Δ*B*_*ij*_ and Δ*B*_*i*_ is maximized (Fig. 5). At *K*≈*L*, Δ*B*_*ij*_ for beneficial becomes negligible, and 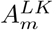 is dominated by the combinatorial factors described in Section 3.

#### 4.1.1 Number of local maxima

In the additive limit, the local maximum is a global maximum and *W* = 1 for all schemes. In the RMF, at 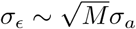, *W* increases dramatically for the beneficial and deleterious scenarios, with *W* ^dele^ exceeding *W* ^bene^ before relaxing to its HoC limit. This transition arises due to an asymmetry in the local maxima: For intermediate *σ*_*ϵ*_, deleterious ascertainment is more likely to produce local maxima at *m* = 2 as compared to beneficial ascertainment at *m* = 1 (Fig. S5). In contrast, in the LK, *W* increases approximately linearly with *K*, with order approximately preserved across ascertainment schemes. As in the RMF model, at intermediate epistasis levels, *K, W* ^dele^ significantly deviates from *W* ^bene^.

Indeed, as in the HoC, *W*_*m*_ differs across schemes. In the additive limit, under random ascertainment, *W*_*m*_ is determined by the number of beneficial mutations available to the ancestral sequence. Assuming deleterious and beneficial mutations are equally likely, 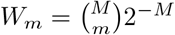, a rescaling of the same quantity for the HoC model. For small *σ*_*ϵ*_, the fully derived sequence is much less likely to be a local maximum under conditional versus trajectory and beneficial ascertainment (Fig. S5).

#### 4.1.2 Number of accessible paths

Unlike with our prior statistics, the additive case produces different values of *P* across ascertainment schemes (Fig. 3G-H). Under random ascertainment, E[*P* ^rand^] = *M* !2^*−M*^, the total number of paths multiplied by the probability that the derived sequence is the global maximum. Trivially, trajectory and beneficial ascertainment result in maximal *P*, while deleterious yields zero accessible paths. *P* ^cond^, however, depends critically on the fitness difference, *F*_*M*_ − *F*_0_, achieved in the uphill adaptive walk (Section S2). Here, *F*_*M*_−*F*_0_ is often small enough such that the additive effects of “conditional” mutations are not exclusively beneficial. Any change to the adaptive dynamics resulting in an increased fitness difference would bring *P* ^cond^ closer to *P* ^traj^ in an additive landscape.

As the amount of epistasis increases, *P* decreases across all ascertainment schemes in both landscapes, with the landscapes yielding qualitatively similar *P* statistics:

For 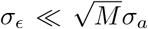, beneficial ascertainment continues to maximize *P* due to the disproportionate contribution of additive effects to phenotypic variance in the CME library (Fig. 4). As *σ*_*ϵ*_ grows, beneficial ascertainment samples mutations with relatively large HoC components, increasing the amount of epistasis in the local landscape. Likewise, for small *K*, the additive effects of beneficial mutations (∝ Δ*B*_*i*_Δ*f*_*i*_) exceed their epistatic contributions to phenotypic variance in the local landscape (∝ Δ*B*_*ij*_; Fig. 5). As *K* grows, however, beneficial mutations share more neighborhoods relative to random mutations (Δ*B*_*ij*_ increases), overwhelming the mutations’ beneficial first order effects, and inducing more rugged, less navigable local landscapes (higher *W* and lower *P* ). As a result, at intermediate amounts of epistasis, *P* ^bene^ dips below that of trajectory and conditional in both landscapes, until reaching its HoC limit of approximately zero.

In the RMF, trajectory ascertainment initially yields more modest *P* relative to beneficial. As with *W*, this result depends critically on the strength of selection. Along a simulated trajectory, mutations are only required to increase fitness relative to the current background, which may be achieved via small or large additive fitness gains, or a large positive HoC component. The trajectory landscape is thus “more rugged” than the beneficial for small *σ*_*ϵ*_ (Fig. 3G). Trajectory ascertainment in the LK landscape qualitatively recapitulates this result, with *P* ^bene^ greater than or equal to *P* ^traj^ until a critical *K* value, where second order effects contribute substantially more variance in beneficial versus trajectory libraries (Fig. 3D).

As *σ*_*ϵ*_ and *K* grow, differences between *P* ^traj^ and *P* ^cond^ shrink (Fig. 3G-H). Fitness gains along the trajectory in the RMF are increasingly mediated by large non-heritable HoC effects, while trajectory mutations in the LK become approximately as connected, or epistatic, as random mutations (Figs. 4 and 5A,C).

**Figure 5.**
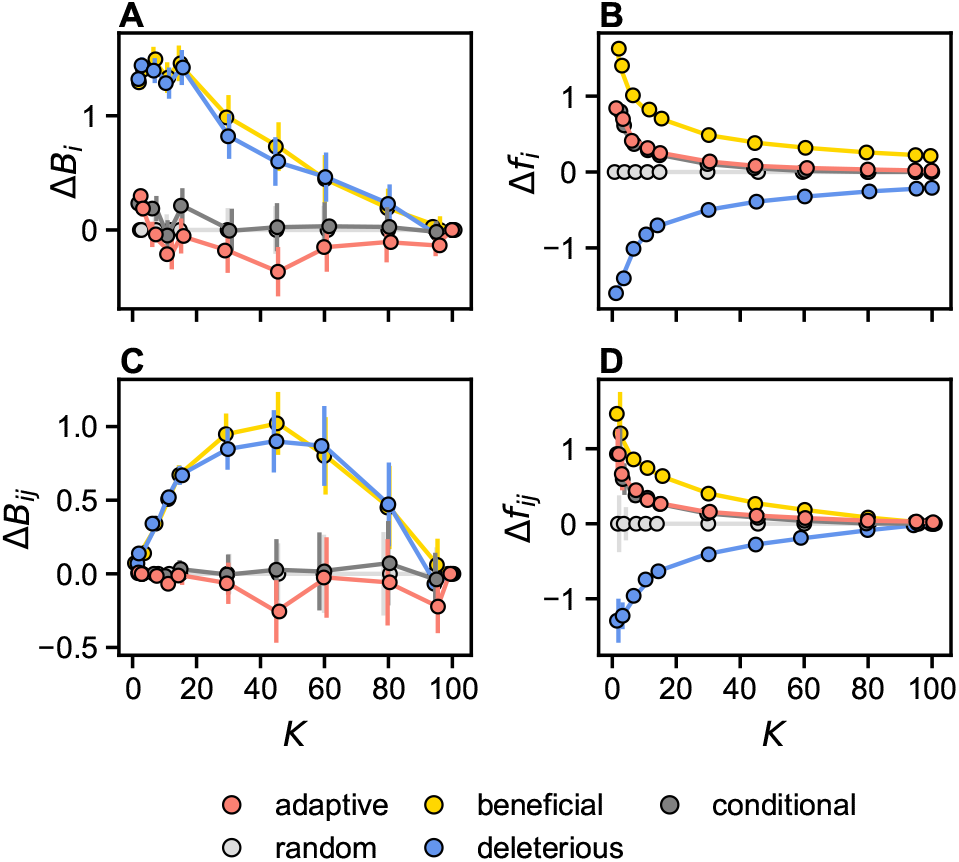
Effects of ascertainment on mutation features in LK landscapes. **(A)** The difference in the average number of neighborhoods that loci from ascertained libraries participate in relative to random loci, 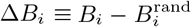, as a function of *K*, with 1 ≤ *K* ≤*L* = 100. **(B)** The average difference in neighborhood contribution attributed to a single mutation,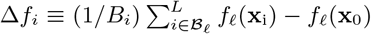, where **x**_i_ and **x**_0_ are the mutant and ancestral sequences, respectively. **(C)** The average difference between number of neighborhoods that loci in each of the beneficial, deleterious, trajectory, and conditional libraries share relative to loci in the random library, 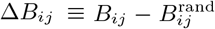. **(D)** The average fitness difference associated with neighborhoods that loci in the same library share, 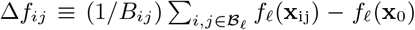, where **x**_*ij*_ is the double mutant. Each point is the average of *N* = 1,000 simulations with *L* = 100 loci in the global landscape. Error bars represent 95% confidence intervals for each statistic.

## 5 Ascertainment biases in empirical datasets

To evaluate whether the theoretical and simulation-based results presented in the previous sections are borne out in empirical datasets, one would ideally experimentally recapitulate the ascertainment scenarios for some set of ancestral sequences. However, as of yet, such an experiment has not been conducted (though see Bank et al., 2016). Therefore, we instead take advantage of a small set of empirical protein fitness landscapes in which the phenotypes of all (or nearly all) possible 20^*M*^ combinations of amino acids at *M* loci have been measured: protein G domain B1 (GB1) binding immunoglobulin (*M* = 4, Wu et al., 2016), and ParD3 binding its cognate and non-cognate toxins, ParE3 and ParE2, respectively (*M* = 3, Lite et al., 2020). It is important to note that these datasets are themselves ascertained. The mutations in GB1 were selected from among the “most epistatic” in an earlier mutagenesis study [Olson et al., 2014, Wu et al., 2016]. The three loci in ParD3 are a subset of residues co-evolving in the ParD-ParE system, at the protein interface, and sufficient to “rewire” the specificity of ParD3 from ParE3 to ParE2 [Aakre et al., 2015, Lite et al., 2020].

We computationally mimic the ascertainment scenarios via biased sub-sampling of the full 20^*M*^ landscapes (Section 7). The three statistics, 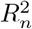, *W*, and *P*, are computed for each scenario and averaged over realizations of the ancestral background.

Sampling from the twenty possible alleles at each site approximates, albeit imperfectly, the *M*≪ *L* regime studied throughout. To see this, let us imagine that one of the empirical landscapes followed the HoC model. In this case, the fitness of each of the 20^*M*^ sequences would be given by an *iid* random normal. Observing many versus two alleles at a given site, however, alters the connectivity of the genotype space: one cannot observe combinations of alleles at the same site. As a result there are 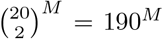 possible biallelic libraries of size 2^*M*^ compared to 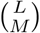 in a biallelic global HoC landscape consisting of 2 ^*L*^ sequences. By this analogy, *M* = 3, 4 for landscapes of size 20^*M*^ correspond to binary, global landscapes with *L* ≈346 and *L* ≈ 422 loci, respectively. Similarly, in the RMF model, each of the 20*M* alleles would have an intrinsic, additive effect, while each of the 20^*M*^ sequences would be associated with an *iid* HoC component. In the LK, the *M* locus neighborhoods, ℬ_*i*_, would consist of a maximum of *M* loci, i.e., 1≤*K*≤*M*, with the 20^*K*^ genotypes associated with a given neighborhood assigned *iid* random variables.

### 5.1 Variance explained by interactions of a given order

In all three datasets, random and deleterious ascertainment exhibit reduced 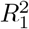 (Fig. 6A; Fig. S11). Deleterious ascertainment tends to select mutations at or below the detection threshold of an experimental assay, where noise dominates fitness differences among mutants [Faure et al., 2020, Carlson et al., 2025]. Thus, it is not surprising that the deleterious scenario closely recapitulates the 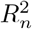 statistics of the HoC model, which can be viewed as a “pure noise” model (Fig. 6A). Random ascertainment likely yields a milder instantiation of this effect as the fitness distributions of all three datasets are enriched for deleterious mutants (Fig. S6). To test this hypothesis, we permuted the genotypes of the samples in each library and recomputed the variance explained by interactions of each order, *Ã*_*m*_. In all datasets, all scenarios excluding deleterious exhibited substantial deviations from the empirical null for *Ã*_1_ (Fig. S7). Deviations at higher orders were less pronounced, indicating that higher 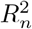 for *n >* 1 in the beneficial, trajectory, and conditional scenarios is primarily achieved via inflation of first-order effects.

**Figure 6.**
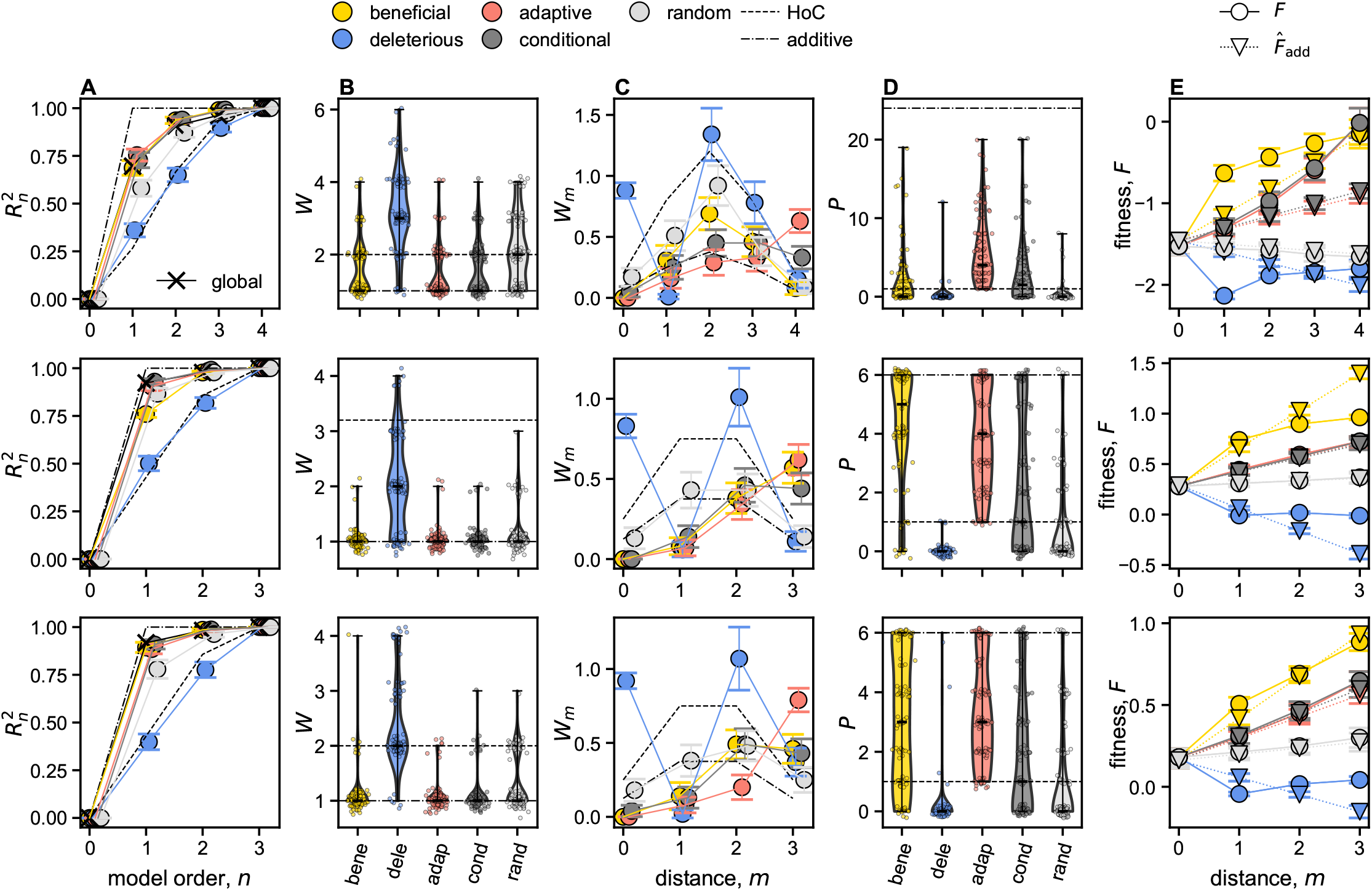
Ascertainment in empirical landscapes. Ascertained mutational libraries were sub-sampled *N* = 100 times from the approximately complete fitness landscape for (first row) *M* = 4 loci in GB1 with respect to binding IgG-Fc [Wu et al., 2016], and *M* = 3 loci in ParD3 with respect to binding (middle row) ParE3 and (bottom row) ParE2. **(A)** Estimates of 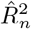as a function of model order *n* for each ascertainment scheme, with analytic expectations of the statistic for House of Cards (HoC) and additive landscapes represented by dashed and dashed-dotted lines, respectively. 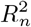 values estimated from the entire dataset are indicated by ×. Error bars represent 95% confidence intervals. **(B)** Violin plots of the total number of local maxima, *W*, with average indicated by the black dash. HoC and additive expectations denoted as in **A. (C)** The number of local maxima as a function of distance from the ancestral sequence, *W*_*m*_, for each ascertainment scheme. All plot features are otherwise as in **A. (D)** Violin plots of the number of uphill accessible paths for each ascertainment scheme, with all plot features otherwise as in **B. (E)** Average observed fitness, *F* (solid line, circles), and predicted fitness based on an additive model fitted with the entire dataset, 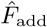 (dotted line, triangles), as a function of distance from the ancestral sequence.

We further hypothesized that systematic deviations from additive predictions, 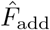 — estimated from the complete datasets — may provide additional insight into the basis of variation in 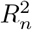 across schemes. In particular, the comparison of observed fitness, *F*, and 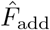 revealed striking differences between beneficial versus trajectory and conditional ascertainment across the datasets. In the ParD3-ParE2 landscape, *F* on average agrees with 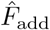 for all three of these ascertained landscapes (Fig. 6E, bottom), consistent with the approximately additive global landscape implied by 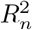 . For ParE3, however, 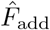 exceeds *F* in the beneficial library for *m* ≥2, consistent with so-called diminishing returns epistasis (Fig. 6E, middle). This phenomenon is most likely due to a global nonlinearity, e.g., induced by an upper measurement threshold [Domingo et al., 2019, Carlson et al., 2025], but could also arise due to an entropic effect in the presence of pervasive specific interactions [Reddy and Desai, 2021]. The presence of a global nonlinearity would also explain the observed reduction in *R*^2^ under beneficial ascertainment in ParE3, as global nonlinearities in the genotype-to-fitness map can inflate estimates of higher-order epistasis if not explicitly modeled [Otwinowski et al., 2018, Domingo et al., 2019, Carlson et al., 2025]. We suspected that this difference between the ParE2 and ParE3 landscapes may result from the fact that ParE3 (and not ParE2) is the cognate toxin of the wildtype sequence. Sequences are thus more likely to reside closer to a fitness optimum (or plateau) for ParE3 versus ParE2, resulting in saturating fitness gains and depressed 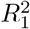 in the former but not the latter.

In contrast, in GB1, average *F* under beneficial ascertainment exceeds additive predictions most substantially for *m* = 1 (Fig. 6E, top). These results are consistent with a highly epistatic landscape: Beneficial ascertainment likely selects mutations that interact well with each of the *M* −1 ancestral alleles. The presence of second and higher-order epistasis, however, renders these initially favorable mutations deleterious with increasing probability as the number of mutations increases, a “regression-to-the-mean” effect. Trajectory ascertainment exhibits the opposite trend, with average *F* exceeding 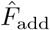 by a larger margin with increasing *m*. By construction, trajectory ascertainment guarantees a fitness increase with *m*, and thus minimizes antagonistic higher-order epistasis.

The parity of conditional with trajectory ascertainment in all datasets suggests that restriction to similarly fit derived sequences is sufficient to ensure similarly fit intermediates, though this similarity in part results from the two scenarios sampling similar alleles (Figs. S8 to S10).

### 5.2 Local maxima and accessible paths

The more “epistatic” deleterious and random landscapes exhibited higher *W* compared to the other three scenarios (Fig. 6B). In addition, *W*_*m*_ for these two scenarios was qualitatively consistent with an underlying HoC landscape (Figs. 2 and 6C).

Interestingly, despite only small differences in 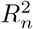, beneficial ascertainment in GB1 exhibited significant differences in *W*_*m*_ from the trajectory and conditional scenarios, even though *W* was similar (Fig. 6B-C). In fact, *W*_*m*_ under beneficial ascertainment resembled that of random with, approximately, 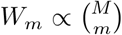, and a constant of proportionality interpolating between the additive and HoC expectations (Fig. 6C). *P* was also significantly lower in the beneficial versus trajectory scenario (Fig. 6D, top), consistent with the observed *W*_*m*_ and the “regression-to-the-mean” effect described above (Fig. 6E, top).

While small differences were observed between *W*_*m*_ for trajectory and conditional scenarios, these were only statistically significant for *m* = 4 in GB1 and *m* = 2, 3 in ParD3-ParE2 (two-sided *t*-test, *p <* .001; Figs. S12 to S14). As discussed, trajectory ascertainment inflates the probability that the derived sequence is a local maximum. In these data-driven simulations, such fitness differences may fall within the experimental margin of error, overstating differences among the schemes. Thus, while *P* in the trajectory scenario exceeded that of conditional in all datasets (two-sided *t*-test, *p <* .005; Figs. S12 to S14), this result may be due primarily to measurement error.

## 6 Discussion

Researchers have previously appreciated the effects of experimental design choices (e.g., genetic background, mutation choice, assayed phenotype, and library size) on the statistics of measured fitness landscapes [Szendro et al., 2013, Bank et al., 2016, Pokusaeva et al., 2019, Agarwala and Fisher, 2019], but have heretofore lacked explicit expectations for the effects of these choices on local landscape statistics.

In the present work, we systematically investigated two key features of CME design: the choice of which mutations and, to a lesser extent, how many mutations *M* to study. We analyzed the consequences of these choices on several oft-used summary statistics in both idealized and empirical fitness landscapes. As expected, we find that the effects of ascertainment depend on the underlying statistics of the larger global landscape. In highly epistatic landscapes, we find that choosing mutations which have occurred along a single adaptive trajectory tends to produce less epistatic local landscapes (i.e., higher 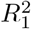) than the other ascertainment scenarios considered. This result is consistent with a prior simulation-based study [Otwinowski and Plotkin, 2014] which demonstrated that sampling from a population under selection results in smoother fitness landscapes. Such sampling was shown to exacerbate biases induced by incomplete sampling of the genotype space and model misspecification (i.e., the omission of true higher-order terms). Our results further build on this work by considering the effects of non-random ascertainment which persist upon complete sampling of the genotype subspace defined by the *M* focal mutations, i.e., when the model is correctly specified.

Our calculations in the HoC landscape demonstrate that these distortions arise from fundamental differences in how the combinatorial library is partitioned with respect to the atypical fitness values. In the beneficial and deleterious scenarios, the “atypical” genotypes are those at a distance of one mutation from the ancestral sequence. In contrast, in the trajectory scenario, each of the atypical genotypes is at a distance of 1, 2, …, *M* mutations from the ancestral sequence. These “assignments” place distinct constraints on the interactions between loci, with pronounced effects in the highly epistatic HoC landscape.

We further show that even when differences in the 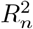 statistics are small, ascertained libraries may exhibit meaningful differences in the number of local maxima *W*, their locations in the landscape *W*_*m*_, and the number of accessible paths *P* . Our analyses of the tunable RMF and LK landscapes show that these effects are highly epistasis-dependent. In near-additive RMF landscapes, beneficial ascertainment produces the most accessible landscapes. As the amount of epistasis increases, however, beneficial ascertainment yields the largest *W* and among the smallest *P* . These patterns are explained by changes in the constitution of beneficial mutations as the amount of epistasis in the global landscape increases. In RMF landscapes, beneficial mutations become relatively more epistatic as the variance of the HoC fitness component increases. In contrast, at small *K* in the LK model, beneficial mutations tend to congregate in neighborhoods with large effects on fitness, resulting in more epistatic local landscapes. More generally, global features of a fitness landscape determine the characteristics of a given mutation type, which in turn, alters the statistics of the local landscape induced by sampling a biased set of mutations.

While not emphasized here, the fitness of the ancestral sequence is also likely to impose substantial constraints on the statistics of a CME. Indeed, a long history of theoretical [Fisher, 1930, Orr, 1998, Blanquart et al., 2014, Reddy and Desai, 2021] and empirical [Kryazhimskiy et al., 2014, Couce et al., 2024] works have characterized systematic changes in the distribution of fitness effects as a function of fitness of the background sequence under different assumptions about the global landscape. The distribution of epistatic effects is also expected to change along an adaptive trajectory [Draghi and Plotkin, 2013, Greene and Crona, 2014, Agarwala and Fisher, 2019]. Thus, considering mutations from distinct parts of longer adaptive walks may yield different results.

While not modeled in our theoretical treatment, empirical fitness estimates are often heteroskedastic. We observed this phenomenon most profoundly in the deleterious libraries generated from three empirical CMEs. More generally, the ability to robustly estimate both *W* and *P* relies on accurate ordering of the fitness values, which is determined by the signal-to-noise ratio and the underlying distribution of fitness differences [Carlson et al., 2025].

In addition, we restricted our theoretical treatment to bi-allelic landscapes. In practice, many systems of interest, e.g., DNA, RNA, and proteins, explore multi-allelic landscapes. The presence of additional alleles may significantly alter evolutionary dynamics [De Visser and Krug, 2014, Bank et al., 2016].

Finally, we did not introduce global nonlinearities in our theoretical analyses, nor explicitly model them in our analyses of empirical datasets (beyond the standard log-transform of the fitness values). The presence of a nonlinearity could alter the 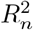 statistic, inflating the contribution of higher-order epistasis to variance explained [Domingo et al., 2019, Carlson et al., 2025]. Indeed, we hypothesized that beneficial ascertainment in the ParD3-ParE3 dataset resulted in diminishing returns epistasis and reduced 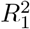. However, absent measurement noise, a global nonlinearity would alter neither *W* nor *P* as these statistics depend only on the ranks of the observed fitness values.

## 7 Materials and Methods

### 7.1 Simulations of random landscapes

In each of the landscape models, an ancestral sequence was sampled uniformly from among all 2^*L*^ sequences in the global landscape. An uphill adaptive walk was simulated by sampling uniformly from among the beneficial mutations at each step. Conditional mutations were sampled via a brute-force search. The other ascertainment schemes were implemented as described in the main text. A simulated replicate was retained only if the uphill walk and conditional search were successful. (See Section S5 for more details.)

### 7.2 Simulations from empirical combinatorial mutagenesis experiments

To mimic the five ascertainment scenarios, local landscapes were sub-sampled from three complete fitness landscapes, for GB1 binding to IgG-Fc [Wu et al., 2016] and ParD3 binding to ParE2 and ParE3, respectively [Lite et al., 2020]. In each simulation, an ancestral sequence was sampled uniformly from all possible sequences. Ascertained libraries of size 2^*M*^ were then generated conditional on the ancestral sequence, according to the five described schemes (see Section S6). A simulation was retained only if fitness was measured for all 2^*M*^ sequences in each ascertained library.

## 8 Code and data availability

All experimental data were previously made available to the public [Wu et al., 2016, Lite et al., 2020, Park et al., 2024]. All of the code required to reproduce the analyses and figures is available at: https://github.com/marync/mutation_choice.

## 9 Acknowledgments

We thank Alasdair Hastewell and attendees of the NITMB research-in-progress for helpful conversations. CB was supported by the U.S. National Science Foundation Graduate Research Fellowship Program under Grant No. DGE-2140743. MOC was supported by a National Institute for Theory and Mathematics in Biology (NITMB) fellowship supported by grants from the NSF (DMS-2235451) and Simons Foundation (MP-TMPS-00005320). MMD was supported by grants R01-GM104239 and R35-GM161612 from the NIH. Generative AI (Claude Opus 4.8 and Sonnet 5.5) was used, in conjunction with manual coding, to write some of the simulation, analysis, and plotting scripts. Claude Opus 5.5 was used to check for and fix (with supervision) typos in the manuscript.

## 10 Author contributions

CB, MMD, and MOC designed the research. CB and MOC performed the research. CB and MOC wrote the paper. CB, MMD, and MOC revised the paper.

