## Supplementary text for "Mutation choice biases the structure of empirical fitness landscapes"

October 21, 2026

### S1 Ascertained summary statistics under the House of Cards model

We provide support for several claims in the main text regarding the effects of ascertainment on the focal summary statistics in the House of Cards (HoC) fitness landscape model.

#### S1.1 Random

##### S1.1.1 Variance explained

Under random ascertainment in the HoC model, the amplitude spectrum is given by the number of interactions of a given order (Neidhart et al., 2013; Agarwala and Fisher, 2019). We derive this result here for completeness. Let $\mathbf{m} \in \{0,1\}^M$ specify an interaction of order $m \in \{1, 2, \ldots, M\}$,

$$\begin{aligned} \mathbb{E}[\beta_{\mathbf{m}}^2] &= \frac{1}{2^{2M}} \sum_{\mathbf{x},\mathbf{x}' \in \mathcal{M}} \mathbb{E}[F(\mathbf{x})F(\mathbf{x}')] \prod_{\ell \in \mathbf{m}} x_\ell x'_\ell \\ &= \frac{1}{2^{2M}} \sum_{\mathbf{x} \in \mathcal{M}} \mathbb{E}[F(\mathbf{x})^2] \\ &= \frac{1}{2^M}, \end{aligned} \tag{S1}$$

and thus,

$$\mathbb{E}[A_m] = \sum_{|\mathbf{m}|=m} \mathbb{E}[\beta_{\mathbf{m}}^2] = \binom{M}{m} \frac{1}{2^M}, \tag{S2}$$

proportional to the number of interactions of order $m$, as claimed in the main text and derived previously (Neidhart et al., 2013).

#### S1.2 Conditional

##### S1.2.1 Variance explained

Under conditional ascertainment in the HoC model, the expected value of the variance explained by a model of order $n$, $R_n^2$, is equal to that under random ascertainment. This result is due to the fact that, as is the case under random ascertainment, $A_m$ is proportional to the number of interactions of order $m$. We solve for $A_m$ under conditional ascertainment,

$$\begin{aligned} \mathbb{E}[\beta_{\mathbf{m}}^2] &= \frac{1}{2^{2M}} \sum_{\mathbf{x},\mathbf{x}' \in \mathcal{M}} \mathbb{E}[F(\mathbf{x})F(\mathbf{x}')] \prod_{\ell \in \mathbf{m}} x_\ell x'_\ell \\ &= \frac{1}{2^{2M}} \sum_{\mathbf{x} \in \mathcal{M}} \mathbb{E}[F(\mathbf{x})^2] \\ &= \frac{1}{2^{2M}} \left[\sigma_g^2 + \mu_g^2 + (2^M - 1)\right], \end{aligned} \tag{S3}$$

where the cross terms cancel as in Eq. S1; the fitness value of the derived sequence is uncorrelated with the fitness values of all intermediate sequences. Thus, as with random ascertainment, $A_m \propto \binom{M}{m}$, the number of interactions of order $m$.

In practice, there will be a positive correlation between the fitness values of the ancestral and derived sequence as an adaptive trajectory will tend to reach a higher fitness value when starting from a high fitness ancestor in the HoC landscape. However, this correlation will contribute only an order $2^{-2M}$ correction.

#### S1.3 Adaptive trajectory scenario

##### S1.3.1 Variance explained

As stated in the main text, interaction terms of the same order will result in distinct partitions of the atypical sequences along an adaptive path. For example, assuming that the mutations are labeled in order of their occurrence along the adaptive trajectory, $\beta_{12}$ partitions the observed adaptive mutants into sets $((1),(2,3,4))$, where adaptive mutant 1 contains the first mutation, 2 the first two mutations, and so on and so forth. In contrast, $\beta_{24}$ partitions the mutants into $((1,4),(2,3))$ as both mutants 1 and 4 contain an even number of flips relative to the ancestral sequence, while 2 and 3 contain an odd number of flips. We thus write the expected value of an interaction of order $m$ specified by the vector $\mathbf{m} \in \{0,1\}^M$ for which $|\mathbf{m}| = m$,

$$\mathbb{E}[\beta_{\mathbf{m}}^2] = \frac{1}{2^{2M}} \left( M(\sigma_g^2 + \mu_g^2) + (2^M - M) + 2\left[ \binom{N_{\mathbf{m}}^+}{2} + \binom{N_{\mathbf{m}}^-}{2} - N_{\mathbf{m}}^+ N_{\mathbf{m}}^- \right] \mu_g^2 \right), \tag{S4}$$

where $N_{\mathbf{m}}^+$ and $N_{\mathbf{m}}^-$ denote the number of adaptive mutants for which the interaction $\mathbf{m}$ contributes positively and negatively, respectively. The amplitude spectrum, $A_m$, is found by summing Eq. S4 over all $\binom{M}{m}$ terms of order $m$ for each $m = 1, \ldots, M$.

In Eq. S4, the expected squared $m^{\text{th}}$ order coefficient is expressed in terms of $N_{\mathbf{m}}^+$ and $N_{\mathbf{m}}^-$, the number of adaptive mutants with net positive and negative sign flips with respect to the $m$ focal loci, respectively. For a given interaction term, this quantity is computed,

$$N_{\mathbf{m}}^+ = \sum_{i=1}^{M} \left[ \left( \sum_{j \in \mathbf{m}} \mathbb{1}\{i \geq j\} \right) \bmod 2 \right], \tag{S5}$$

with $N_{\mathbf{m}}^- = M - N_{\mathbf{m}}^+$. The inner sum is over all loci $j$ involved in the interaction, i.e., $m_j = 1$. If $i < j$ then the $i^{\text{th}}$ adaptive mutant will be in the ancestral state at locus $j$. If $i \geq j$ then the $i^{\text{th}}$ adaptive mutant will be in the opposite state as ancestor at locus $j$. When the inner sum is even, the $i^{\text{th}}$ mutant exhibits the same sign with respect to the focal loci as the ancestor, i.e.,

$$\prod_{j \in \mathbf{m}} x_j^{(i)} = \prod_{j \in \mathbf{m}} x_j^{(0)},$$

where $x_j^{(0)}$ and $x_j^{(i)}$ are the genotypes of the ancestor and the $i^{\text{th}}$ adaptive mutant at locus $j$, respectively.

##### S1.3.2 Number of local maxima

As discussed in the main text, the number of local maxima, $W^{\text{traj}}$, in the trajectory landscape is trivially reduced due to the fact that mutants along an adaptive trajectory are by definition not maxima. In addition, trajectory ascertainment will also affect the probability that non-adaptive mutants are local maxima. To find the expected number of local maxima in the trajectory landscape, we consider the probability that non-trajectory mutants at a given distance $d$ from the ancestor are local maxima. For a library of size $2^M$

and $1 \leq d \leq M-1$, and conditioning on the trajectory, $\mathcal{T}_M$,

$$\begin{aligned}\mathbb{E}[W_d|\mathcal{T}_M] = &\left[\binom{M}{d} - M\right]\frac{1}{M+1} + (M-d-1)\left(1-\Phi[F_{(d-1)}]\right)\int_{f=F_{(d-1)}}^{\infty}\Phi[f]^{M-1}\tilde{\phi}[f]df + \\ &(d-1)(1-\Phi[F_{(d+1)}])\int_{f=F_{(d+1)}}^{\infty}\Phi[f]^{M-1}\tilde{\phi}[f]df + (1-\Phi[F_{(d+1)}])\int_{f=F_{(d+1)}}^{\infty}\Phi[f]^{M-2}\tilde{\phi}[f]df \\ = &\frac{\binom{M}{d}-M}{M+1} + \frac{M-d-1}{M}\left(1-\Phi[F_{(d-1)}]^M\right) + \\ &\frac{d-1}{M}(1-\Phi[F_{(d+1)}]^M) + \frac{1}{M-1}(1-\Phi[F_{(d+1)}]^{M-1}),\end{aligned} \tag{S6}$$

where $\Phi[\cdot]$ is the cumulative distribution function (*cdf*) of a standard normal random variable, and, in an abuse of notation, $\tilde{\phi}[\cdot]$ is the distribution of a standard normal random variable truncated at the appropriate value of $F$. The four terms in Eq. S6 correspond to the mutants at a distance $d$ from the ancestral sequence which (1) have no trajectory neighbors, (2) are neighbors of the $\mathbf{x}_{(d-1)}$, the $(d-1)$-*th* trajectory mutant, (3) are neighbors of the $(d+1)$-*th* trajectory mutant, or (4) are neighbors of both the $(d-1)$-*th* and $(d+1)$-*th* mutant. Using Eq. S6 and typical values of mutants along an adaptive trajectory, we can approximate the expectation of $W^{\text{traj}}$ as,

$$\mathbb{E}[W^{\text{traj}}] \approx \sum_{d=1}^{M-1}\mathbb{E}[W_d|\mathcal{T}_M] + 1 \tag{S7}$$

where we assume that $F_{(i)} \in \mathcal{T}_M$ represent the typical trajectory and that $F_{(0)} \sim \mathcal{N}(0,1)$. To estimate the number of local maxima, we compute Eq. S7 using the average fitness values of mutants sampled along $N = 100$ simulated adaptive trajectories.

##### S1.3.3 Pathway accessibility

By definition, the adaptive path is an accessible path. However, as discussed in the main text, trajectory ascertainment additionally perturbs the number of accessible paths, $P$. Consider a path $p$ which involves the $i^{\text{th}}$ and $j^{\text{th}}$ adaptive mutants, $F_i$ and $F_j$, in a trajectory of length $M+1$ mutations, but no other adaptive mutants. The probability that $p$ is accessible is given by,

$$\mathbb{P}\{p \in \mathcal{A}\} = \frac{(\Phi[F_{(i)}]-\Phi[F_{(0)}])^{i-1}}{(i-1)!} \cdot \frac{(\Phi[F_{(j)}]-\Phi[F_{(i)}])^{j-i-1}}{(j-i-1)!} \cdot \frac{(\Phi[F_{(M)}]-\Phi[F_{(j)}])^{M-j-1}}{(M-j-1)!}, \tag{S8}$$

where each term in Eq. S8 is the probability of correctly ordering subsets of $p$, with the partitioning imposed by $i$ and $j$, the positioning of the trajectory mutants along the path. By enumerating the number of times a given subset of trajectory mutants is included in a path from the ancestral to derived sequence, and under assumptions about the fitness values of the trajectory mutants, we can compute the expected number of accessible paths,

$$\mathbb{E}[P] = \sum_{s \in \mathcal{S}_M}(\#\text{ paths which include } s)\mathbb{P}\{p(s) \in \mathcal{A}\} + (\#\text{ paths in } \emptyset)\mathbb{P}\{p \in \mathcal{A}\}, \tag{S9}$$

where $S_M$ denotes the set of all possible subsets of the trajectory mutants, i.e., $S_M$ is the powerset of $\{1,\ldots,M-1\}$, excluding the empty set (see Fig. S2). To estimate the number of accessible paths, we compute Eq. S8 using the average fitness values of mutants sampled along $N = 100$ simulated adaptive trajectories.

#### S1.4 Beneficial scenario

##### S1.4.1 Variance explained

We first provide a full derivation of the amplitude spectrum, $A_m$, under the approximate beneficial scenario in which the fitness of each of the $M$ single mutants is sampled from a distinct distribution $\mathcal{G}$, with mean $\mu_g$ and variance $\sigma_g^2$. Here, the combinatorial library consists of $2^M - M$ mutants with fitness values sampled from $\mathcal{F}$, a standard normal distribution, and $M$ mutants with fitness values sampled *iid* from $\mathcal{G}$. Under this assumption, we can derive the expected amplitude spectrum, where the expectation is taken over realizations of the landscape. The expectation of the squared sample average, $A_0$, is given by

$$\begin{aligned} A_0 \equiv \mathbb{E}[\mu^2] &= \frac{1}{2^{2M}} \sum_{\mathbf{x},\mathbf{x}' \in \mathcal{M}} \mathbb{E}[F(\mathbf{x})F(\mathbf{x}')] \\ &= \frac{1}{2^{2M}} \left[ M(\sigma_g^2 + \mu_g^2) + (2^M - M) + 2\binom{M}{2}\mu_g^2 \right]. \end{aligned} \tag{S10}$$

The sample variance is defined as

$$Var[\mathbf{F}] := \frac{1}{2^M} \sum_{x \in \mathcal{M}} \left( F(\mathbf{x}) - \bar{F} \right)^2, \tag{S11}$$

with expectation,

$$\mathbb{E}[Var[\mathbf{F}]] = \frac{1}{2^M} \left[ \left[ M(\sigma_g^2 + \mu_g^2) + (2^M - M) \right] \left( 1 - \frac{1}{2^M} \right) - M(M-1)\mu_g^2 \left( \frac{1}{2^M} \right) \right], \tag{S12}$$

compared to $\mathbb{E}[Var[\mathbf{F}]] = 2^{-M}(2^M - 1)$ under random choice of mutations, i.e., when all fitness values are *iid* standard normal. We solve for each term in Eq. 1 of the main text (defined for a sub-landscape of size $2^M$),

$$\mathbb{E}[\mu] = \frac{1}{2^M} \sum_{\mathbf{x} \in \mathcal{M}} F(\mathbf{x}) = \frac{M}{2^M} \cdot \mu_g, \tag{S13}$$

where $\mu_g$ is the expected value of $F_i \sim \mathcal{G}$. Similarly,

$$\mathbb{E}[\beta_\ell] = \frac{1}{2^M} \left( \sum_{x_\ell = 1} \mathbb{E}[F(\mathbf{x})] - \sum_{x'_\ell = -1} \mathbb{E}[F(\mathbf{x}')] \right) = 0, \tag{S14}$$

as each of the $M$ mutations is equally likely to be in either state ($x_\ell \in \{-1, 1\}$) in the ancestor. We find the squared term similarly,

$$\begin{aligned} \mathbb{E}[\beta_\ell^2 | \mathbf{x}_{\text{anc}}] &= \frac{1}{2^{2M}} \left( \sum_{x_\ell=1} \sum_{x'_\ell=1} \mathbb{E}[F(\mathbf{x})F(\mathbf{x}')] + \sum_{x_\ell=-1} \sum_{x'_\ell=-1} \mathbb{E}[F(\mathbf{x}')F(\mathbf{x})] - 2 \sum_{x_\ell=1} \sum_{x'_\ell=-1} \mathbb{E}[F(\mathbf{x}')F(\mathbf{x})] \right) \\ &= \frac{1}{2^{2M}} \left( M(\sigma_g^2 + \mu_g^2) + 2^M - M + 2\binom{M-1}{2}\mu_g^2 - 2(M-1)\mu_g^2 \right) \\ &= \frac{1}{2^{2M}} \left( M\sigma_g^2 + (2^M - M) + \mu_g^2(M + (M-1)(M-2) - 2(M-1)) \right) \\ &= \frac{1}{2^{2M}} \left( M\sigma_g^2 + (2^M - M) + (M^2 - 4M + 4)\mu_g^2 \right) \end{aligned} \tag{S15}$$

where, regardless of the ancestral sequence, each site $\ell$ partitions the set of *beneficial* variants into two sets of size 1 and $M - 1$.

By the same symmetry argument, we have that the expected value of each higher order coefficient is also zero. We now solve for the expected square of $\beta_{\ell m}$,

$$
\begin{aligned}
\mathbb{E}[\beta_{\ell m}^2] &= \frac{1}{2^{2M}}\left(\sum_{x'_\ell x_m=1}\mathbb{E}[F(\mathbf{x})F(\mathbf{x}')] + \sum_{x'_\ell x_m=-1}\mathbb{E}[F(\mathbf{x}')F(\mathbf{x})] - 2\sum_{x_m x_\ell=-1}\sum_{x'_m x'_{\ell m}=1}\mathbb{E}[F(\mathbf{x}')F(\mathbf{x})]\right)\\
&= \frac{1}{2^{2M}}\Big[2(\sigma_g^2+\mu_g^2) + \left(\frac{2^M}{2}-2\right)\sigma^2 + 2\mu_g^2\\
&\qquad + (M-2)(\sigma_g^2+\mu_g^2) + \left(\frac{2^M}{2}-(M-2)\right)\sigma^2 + 2\binom{M-2}{2}\mu_g^2 - 2\left(2(M-2)\mu_g^2\right)\Big]\\
&= \frac{1}{2^{2M}}\Big[M(\sigma_g^2+\mu_g^2) + (2^M-M)\sigma^2 + \left[2+2\binom{M-2}{2} - 4(M-2)\right]\mu_g^2\Big],
\end{aligned}
\tag{S16}
$$

from which we can generalize to arbitrary order $m \leq M$,

$$
\mathbb{E}[\beta_{(m)}^2] = \frac{1}{2^{2M}}\left(M(\sigma_g^2+\mu_g^2) + (2^M-M)\sigma^2 + 2\left[\binom{m}{2} + \binom{M-m}{2} - m(M-m)\right]\mu_g^2\right). \tag{S17}
$$

This result follows from the fact that each atypical mutant only "flips" one of the ancestral states. Each interaction of order $m$ involves $m$ positions, of which a given atypical sequence will be identical to the ancestral sequence or have one flipped state. Consider the third order interaction pertaining to loci $\ell$, $m$, and $n$. This interaction will contribute $\beta_{\ell mn}x_\ell(-x_m)x_n = -\beta_{\ell mn}x_\ell x_m x_n$ to the fitness of the single mutant at site $m$, and likewise for single mutants $\ell$ and $n$, where $x_i$ is the genotype of the ancestor at position $i$. However, the contribution will be $\beta_{\ell mn}x_\ell x_m x_n$ for all other single mutants. Thus, an interaction of order $m$ partitions the atypical, beneficial mutants into sets of size $m$ and $M-m$.

##### S1.4.2 Better approximations to the beneficial scenario

Above, and in the main text, we simplified the beneficial scenario for ease of exposition and to guide intuition. We improve upon the approximation to $R_n^2$ in the main text by more precisely modeling the distributions of the beneficial mutants.

In the beneficial scenario, the fitness values of the single mutants are the top $M$ order statistics of $L$ draws from a standard normal distribution. Following Blom (Eq. 6.10.3, Blom 1958), the expected value of the $m^{\text{th}}$ largest order statistic is well approximated by

$$
\mu_{(m)} \equiv \mathbb{E}[F_{(m)}] \approx \Phi^{-1}(p_m), \quad \text{where} \quad p_m = \frac{L-m+1-\alpha}{L-2\alpha+1}, \tag{S18}
$$

where $\Phi^{-1}(\cdot)$ is the inverse cumulative distribution function (*cdf*), or quantile function, of the standard normal distribution, and $\alpha = 3/8$. The variance of the $m^{\text{th}}$ order statistic can be approximated using the Delta method,

$$
\sigma_{(m)}^2 \equiv \mathbb{V}[F_{(m)}] \approx \frac{p_m(1-p_m)}{(L+2)\left[\phi(\Phi^{-1}(p_m))\right]^2}, \tag{S19}
$$

where $\phi(\cdot)$ is the probability density function of the standard normal distribution and $p_m$ is defined as in the previous equation (Eq. 4.6.4 of David and Nagaraja, 2003).

The interaction terms are no longer *iid* due to the fact that each of the single mutants has a distinct distribution. Without loss of generality, assume that the beneficial mutants are ordered by fitness, such that $F_{(1)} \geq \ldots \geq F_{(M)}$. The expected square of a given interaction term is then,

$$
\mathbb{E}[\beta_{\mathbf{m}}^2] \approx \frac{1}{2^{2M}}\left[\sum_{i=1}^{M}(\sigma_{(i)}^2+\mu_{(i)}^2) + (2^M-M)\sigma^2 + 2\left(\sum_{\substack{i,j\in\mathbf{m}\\ j<i}}\mu_{(i)}\mu_{(j)} + \sum_{\substack{i,j\notin\mathbf{m}\\ j<i}}\mu_{(i)}\mu_{(j)} - \sum_{i\in\mathbf{m}}\sum_{j\notin\mathbf{m}}\mu_{(i)}\mu_{(j)}\right)\right], \tag{S20}
$$

where $\mathbf{m}$ is an $m$-length vector indicating which sites participate in the interaction and $\mu_{(i)}$ and $\sigma^2_{(i)}$ are approximations to the mean and variance of the $i^{\text{th}}$ beneficial mutant (Eq. S18 and S19). Eq. S20 additionally ignores correlations among the order statistics. More accurate approximations could thus be obtained by approximating the covariances among pairs of order statistics (see Chapter 4 of David and Nagaraja, 2003). The amplitude spectrum, $A_m$, is then found by summing over all possible combinations of $m$ loci for each $m = 1, \ldots, M$.

#### S1.5 A microscopic view of the HoC model

The fitness of a sequence in the HoC landscape can be represented as,

$$F(\mathbf{x}) = \mu + \sum_{\ell=1}^{L} \beta_\ell x_\ell + \sum_{\ell<m} \beta_{\ell m} x_\ell x_m + \ldots + \sum_{|\mathbf{m}|=L} \beta_{\mathbf{m}} \prod_{\ell \in \mathbf{m}} x_\ell, \tag{S21}$$

with interactions up to order $L$, the length of the sequence, and where $\beta_\cdot \overset{iid}{\sim} \mathcal{N}(0, 2^{-L})$ such that $F(\cdot) \overset{iid}{\sim} \mathcal{N}(0,1)$. Sub-sampling to a combinatorial library of size $2^M$ necessarily constrains sequences in the library to the $M$-dimensional hypercube,

$$\tilde{F}(\mathbf{x}) = \tilde{\mu} + \sum_{m=1}^{M} \tilde{\beta}_m x_m + \sum_{\ell<m} \tilde{\beta}_{\ell m} x_\ell x_m + \ldots + \sum_{|\mathbf{m}|=M} \tilde{\beta}_{\mathbf{m}} \prod_{\ell \in \mathbf{m}} x_\ell, \tag{S22}$$

where $\tilde{\mu}$ absorbs the effects of all first order effects and interactions pertaining to the $L - M$ fixed sites in the library; $\tilde{\beta}_m$ analogously absorbs the effects of all interactions of locus $m$ with the $L - M$ fixed sites; so on and so forth for higher order interactions defined with respect to the combinatorial library. For example,

$$\tilde{\mu} = \mu + \sum_{\ell \notin \mathcal{M}} \beta_\ell x_\ell + \sum_{\ell < m \notin \mathcal{M}} \beta_{\ell m} x_\ell x_m + \ldots,$$

and, for $m, m' \in \mathcal{M}$,

$$\tilde{\beta}_{mm'} = \beta_{mm'} + \sum_{\ell \notin \mathcal{M}} \beta_{mm'\ell} x_\ell + \sum_{n < \ell \notin \mathcal{M}} \beta_{mm'\ell n} x_\ell x_n + \ldots, \tag{S23}$$

where $x_\ell$ is the ancestral state at locus $\ell \notin \mathcal{M}$, and the summations continue up to order $L$ interactions.

Without loss of generality, let us fix the ancestral sequence $\mathbf{x}_0 = \begin{pmatrix} -1 & -1 & \ldots & -1 \end{pmatrix}$. (Fixing the ancestral sequence is akin to considering $\tilde{\beta}_i x_i$ and $\tilde{\beta}_{ij} x_i x_j$, and so on and so forth.) This "reference" encoding will be more useful for understanding the effects of ascertainment on $A_m$. As this choice does not alter the magnitudes of the coefficients, only their signs, it will not affect estimation of $A_m$.

Let us consider the case where $M = 2$. We have

$$\begin{aligned} F_0 &= \tilde{\mu} - \tilde{\beta}_1 - \tilde{\beta}_2 + \tilde{\beta}_{12} \\ F_1 &= \tilde{\mu} + \tilde{\beta}_1 - \tilde{\beta}_2 - \tilde{\beta}_{12} \\ F_2 &= \tilde{\mu} - \tilde{\beta}_1 + \tilde{\beta}_2 - \tilde{\beta}_{12} \\ F_{12} &= \tilde{\mu} + \tilde{\beta}_1 + \tilde{\beta}_2 + \tilde{\beta}_{12}, \end{aligned} \tag{S24}$$

where $F_0$, $F_1$, $F_2$, and $F_{12}$, are the fitness values of the ancestor, the single mutants at loci one and two, and the double mutant, respectively.

##### S1.5.1 Beneficial ascertainment

Under the HoC model with beneficial ascertainment, $F_0 \sim F_{12}$, with $\mathbb{E}[F_0] = \mathbb{E}[F_{12}] = 0$, while $\mathbb{E}[F_1] \approx \mathbb{E}[F_2] \equiv \mu_g > 0$ (and $\mu_g > \tilde{\mu}$). These constraints are satisfied if $\tilde{\beta}_{12} < 0$, $\tilde{\beta}_1 + \tilde{\beta}_2 = 0$ (or $\mathbb{E}[\tilde{\beta}_1] = \mathbb{E}[\tilde{\beta}_2] = 0$), and $\tilde{\mu} + \tilde{\beta}_{12} = 0$. The latter two constraints imply that, unlike in the full (or randomly ascertained) HoC landscape, $\tilde{\beta}_1$ and $\tilde{\beta}_2$, and $\tilde{\mu}$ and $\tilde{\beta}_{12}$ are negatively correlated, respectively. Altogether, these results imply that a library constructed from $M = 2$ beneficial mutants will be enriched for second-order, negative epistasis. This result translates the "regression to the mean" phenomenon—whereby the combination of mutations with extreme effects results in a less extreme phenotype—to its representation in the microscopic Fourier basis.

As $M$ increases, the amplitude spectrum shifts, changing the apportionment of epistasis among different orders (Eq. 8). This apportionment is determined by the number of beneficial mutations in which combinations of loci occur in the same configuration (i.e., $-1$ or 1) relative to the number of pairs in distinct configurations; the partitioning encoded in Eq. 8 of the main text. Together, these results imply that in highly epistatic landscapes such as the HoC, the apparent interactions among loci are highly dependent on the ensemble of mutations and the number of mutations considered. Already, in Eq. S23, we see that the effective interaction term is implicitly conditioned on the ancestral background. Beneficial ascertainment, in turn, implies that the interactions of a single locus $m \in \mathcal{M}$ or two loci $m, m' \in \mathcal{M}$ with the genetic background may be highly atypical.

###### S1.5.2 Trajectory ascertainment

Trajectory ascertainment imposes distinct constraints on the fitness values and thus coefficients of an empirical fitness landscape. For $M = 2$, to a first approximation, we assume that $F_1 \sim F_{12} \sim \mathcal{N}(\mu_g, \sigma_g^2)$ while $F_0 \sim F_2 \sim \mathcal{N}(0, 1)$, where the subscript denotes the mutated alleles with respect to the ancestral genotype. Here, both adaptive mutants are in the same state at the first locus to mutate in the adaptive trajectory, implying that $\mathbb{E}[\tilde{\beta}_1] > 0$ under the reference encoding, where we have indexed the mutations by their order in the trajectory. The distribution of first order coefficients is thus bi-modal,

$$\tilde{\mu} \sim \mathcal{N}\left(\frac{\mu_g}{2}, \frac{\sigma_g^2 + 1}{8}\right), \quad \tilde{\beta}_1 \sim \mathcal{N}\left(\frac{\mu_g}{2}, \frac{\sigma_g^2 + 1}{8}\right), \quad \tilde{\beta}_2 \sim \tilde{\beta}_{12} \sim \mathcal{N}\left(0, \frac{\sigma_g^2 + 1}{8}\right). \tag{S25}$$

As $M$ increases, the distributions of coefficients at each order become mixtures of an increasing number of Gaussians, with means determined by the number of beneficial mutants in which the focal loci appear in the same configuration in the beneficial mutants. Hence, we arrive at Eq. S4.

### S2 Ascertained summary statistics under the additive model

In an additive landscape constructed from $M$ loci, there are either $M!$ or zero uphill paths.

#### S2.1 Random ascertainment

The probability that the $M$ selected loci are in the negative orientation in the ancestral sequence is simply $2^{-M}$. Thus the expected number of paths is $2^{-M}M!$.

#### S2.2 Conditional ascertainment

Let $F_0$ and $F_M$ be the ancestral and derived fitness values from a given adaptive trajectory. The probability that a given path in a library constructed under conditional ascertainment is uphill is determined by the conditional distribution of effect sizes, $s_\ell = -2\lambda_\ell x_\ell$, where $x_\ell$ is the state of the ancestral sequence,

$$\begin{bmatrix} s_1 \\ s_2 \\ \vdots \\ s_M \\ F_M - F_0 \end{bmatrix} \sim \mathcal{N}\left(\mathbf{0}, \begin{bmatrix} 4\sigma_a^2 & 0 & \cdots & 4\sigma_a^2 \\ & 4\sigma_a^2 & 0\cdots & 4\sigma_a^2 \\ & & \ddots & \vdots \\ & & 4\sigma_a^2 & 4\sigma_a^2 \\ & & & 4M\sigma_a^2 \end{bmatrix}\right), \tag{S26}$$

which implies,

$$s_1, \cdots, s_M | F_M - F_0 \sim \mathcal{N}\left(\frac{1}{M}(F_M - F_0)\mathbf{1}, 4\sigma_a^2(I - \frac{1}{M}\mathbf{1}\mathbf{1}^\top)\right), \tag{S27}$$

where the covariance matrix is degenerate, and $s_\ell$ now live on the hyperplane with $\sum s_\ell = F_M - F_0$. To sample $s_\ell$, we let $s_\ell = (F_M - F_0)/M + 2\sigma_a(Z_\ell - \bar{Z})$, where $Z_\ell \overset{iid}{\sim} \mathcal{N}(0,1)$ and $\bar{Z} = \frac{1}{M}\sum_\ell Z_\ell$. It is easily shown that the $s_\ell$ have the correct distribution. If we assume that the typical value of $F_M - F_0 \approx 2M\sqrt{2/\pi}$, then for $M = 4$ and $\sigma_a^2 = 1$, the probability that all effects are positive is approximately 0.36, yielding an expected number of paths of 8.64.

### S3 Ascertained summary statistics in the Rough Mount Fuji model

In the next sections, we consider the additive and House of Cards (HoC) contributions to the fitness values of mutants obtained under distinct ascertainment schemes. This apportionment determines the extent to which the ascertained loci participate in epistatic interactions.

#### S3.1 Variance explained under random ascertainment

For completeness, we derive the approximation for the variance explained by a model of order $n$, $R_n^2$, under random ascertainment in the RMF model presented in the main text (Eq. 15) and derived previously by Neidhart et al. (2013). We begin with the expectation of the square of the first order effect of locus $\ell$,

$$\begin{aligned}
\mathbb{E}[\beta_\ell^2] &= \frac{1}{2^{2M}}\left[\sum_{\mathbf{x}}\sum_{\mathbf{x}'} F(\mathbf{x})F(\mathbf{x}')x_\ell x'_\ell\right] \\
&= \frac{1}{2^{2M}}\left[\sum_{\mathbf{x}}\sum_{\mathbf{x}'}\left(\sum_m \lambda_m x_m + \epsilon\right)\left(\sum_n \lambda_n x_n + \epsilon'\right)x_\ell x'_\ell\right] \\
&= \frac{1}{2^{2M}}\left[\sum_{\mathbf{x}}\sum_{\mathbf{x}'}\mathbb{E}[\lambda_\ell^2(x_\ell x'_\ell)^2] + 2^M\sigma_\epsilon^2\right] \\
&= \sigma_a^2 + \frac{1}{2^M}\sigma_\epsilon^2,
\end{aligned} \tag{S28}$$

and thus, the first element of the amplitude spectrum, $A_1$, is given by,

$$\mathbb{E}[A_1] = M\sigma_a^2 + \frac{M}{2^M}\sigma_\epsilon^2. \tag{S29}$$

The first order effects will only contribute to $A_1$, and thus, the $A_m$ for $m > 1$ are equal to those of the HoC model, i.e., $A_m = \binom{M}{m}2^{-M}\sigma_\epsilon^2$. Combining these results yields the approximate expression for $R_n^2$ provided in the main text (Eq. 15; Fig. S3).

#### S3.2 Distributions of additive and HoC components

The normality of the additive and HoC components of the RMF model enable us to solve explicitly for their contributions to the fitness effect of a given mutation $\ell$, $s_\ell \equiv F_\ell - F$, in an ancestral background $F$. The fitness effect of site $\ell$ can be expressed as,

$$s_\ell = F_\ell - F = F - 2\lambda_\ell x_\ell + \epsilon_\ell - \epsilon - F = -2\lambda_\ell x_\ell + \epsilon_\ell - \epsilon, \tag{S30}$$

where $x_\ell \in \{-1, 1\}$ is the state of the ancestral sequence $\mathbf{x}$ at locus $\ell$; and, $\epsilon$ and $\epsilon_\ell$ are the HoC contributions to the fitness of the ancestor, $F$, and that of the single mutant, $F_\ell$, respectively. Thus,

$$\begin{pmatrix} -2\lambda_\ell x_\ell \\ \epsilon_\ell \\ s \\ F \end{pmatrix} \sim \mathcal{N}\left(\vec{\mathbf{0}}, \begin{pmatrix} 4\sigma_a^2 & 0 & 4\sigma_a^2 & -2\sigma_a^2 \\ & \sigma_\epsilon^2 & \sigma_\epsilon^2 & 0 \\ & & 4\sigma_a^2 + 2\sigma_\epsilon^2 & -2\sigma_a^2 - \sigma_\epsilon^2 \\ & & & L\sigma_a^2 + \sigma_\epsilon^2 \end{pmatrix}\right), \tag{S31}$$

where $\sigma_a^2$ and $\sigma_\epsilon^2$ are the additive and HoC variances, as defined above. The distribution of the additive and HoC contributions conditional on $s$ and $F$ is thus bivariate normal, with means given by

$$\mathbb{E}[-2\lambda_\ell x_\ell | s, F] = \left(\frac{2\sigma_a^2}{2\sigma_a^2 + \sigma_\epsilon^2}\right) s \tag{S32}$$

and

$$\mathbb{E}[\epsilon_\ell | s, F] = \frac{\sigma_\epsilon^2 \left[(\sigma_\epsilon^2 + 2\sigma_a^2)F + (\sigma_\epsilon^2 + L\sigma_a^2)s\right]}{(2\sigma_a^2 + \sigma_\epsilon^2)(\sigma_\epsilon^2 + 2\sigma_a^2(L-1))}, \tag{S33}$$

where the denominator in Eq. S33 is the determinant of the $2 \times 2$ sub-matrix of Eq. S31 specifying the covariance of $s$ and $F$. These equations imply that for a given $F$, $s$, and $\sigma_a^2$, we can compute the precise value of $\sigma_\epsilon^2$ where the HoC component (in expectation) comes to dominate the fitness effect of locus $\ell$. These quantities are also, of course, negatively correlated,

$$\mathbb{C}[-2\lambda_\ell x_\ell, \epsilon_\ell | s, F] = -\frac{2\sigma_a^2\sigma_\epsilon^2}{2\sigma_a^2 + \sigma_\epsilon^2}. \tag{S34}$$

The expected additive effect is proportional to $s$, with constant of proportionality determined by the variance fraction contributed by additive variation to the fitness effect (Eq. S32). Moreover, the expected additive effect is not dependent on $F$ when conditioned on $s$. In contrast, the expected HoC component increases with both $F$ and $s$ (Eq. S33). For large $L$, the conditioning on $s$ dominates the expected value of $\epsilon_\ell$, which becomes proportional to the variance fraction of the HoC component,

$$\mathbb{E}[\epsilon_\ell | s, F] \to \frac{1}{2}\left(\frac{\sigma_\epsilon^2}{2\sigma_a^2 + \sigma_\epsilon^2}\right) s,$$

as gleaned from Eq. S32.

We use the fact that the distribution of $s$ conditional on $F$, i.e., the conditional distribution of fitness effects (dfe), is normally distributed to find an approximation for its maximum. We have

$$s|F \sim \mathcal{N}\left(-\frac{2\sigma_a^2 + \sigma_\epsilon^2}{L\sigma_a^2 + \sigma_\epsilon^2}F, \frac{(2\sigma_a^2 + \sigma_\epsilon^2)(\sigma_\epsilon^2 + 2\sigma_a^2(L-1))}{L\sigma_a^2 + \sigma_\epsilon^2}\right). \tag{S35}$$

Letting $\mu_s = \frac{2\sigma_a^2 + \sigma_\epsilon^2}{L\sigma_a^2 + \sigma_\epsilon^2}$, we see that $\mathbb{E}[s|F] = -\mu_s F$, while $\mathbb{V}[s|F] = \mu_s[2\sigma_a^2(L-1) + \sigma_\epsilon^2]$; both the expected value and variance scale linearly with the amount of variance explained by a single mutation relative to total phenotypic variance. For large $L$, $\mathbb{V}[s|F] \approx 2(2\sigma_a^2 + \sigma_\epsilon^2)$, proportional to the unconditional variance of $s$. A conditional fitness effect thus has a maximum of approximately,

$$\mathbb{E}[s_{\max}|F] \approx -\frac{2\sigma_a^2 + \sigma_\epsilon^2}{L\sigma_a^2 + \sigma_\epsilon^2}F + \sqrt{\frac{(2\sigma_a^2 + \sigma_\epsilon^2)(\sigma_\epsilon^2 + 2\sigma_a^2(L-1))}{L\sigma_a^2 + \sigma_\epsilon^2}}\sqrt{2\ln L} \approx 2\sqrt{(2\sigma_a^2 + \sigma_\epsilon^2)\ln L}, \tag{S36}$$

where the latter approximation follows for large $L$ and $\sigma_\epsilon \ll L\sigma_a^2$. The first term in Eq. S36 reflects the fact that as $F$ increases, the maximum fitness gain decreases, a manifestation of the so-called diminishing returns epistasis effect.

#### S3.3 Fitness components under conditional ascertainment

Let $X_M = -2\sum_{\ell \in \mathcal{M}} \lambda_\ell x_\ell$, be the difference in fitness between the ancestral and derived genotypes due to additive effects, and $\epsilon_M$ the contribution of the HoC component,

$$\begin{pmatrix} X_M \\ \epsilon_M \\ F_M \\ F_0 \end{pmatrix} \sim \mathcal{N}\left(\vec{\mathbf{0}}, \begin{pmatrix} 4M\sigma_a^2 & 0 & 2M\sigma_a^2 & -2M\sigma_a^2 \\ & \sigma_\epsilon^2 & \sigma_\epsilon^2 & 0 \\ & & L\sigma_a^2 + \sigma_\epsilon^2 & (L-2M)\sigma_a^2 \\ & & & L\sigma_a^2 + \sigma_\epsilon^2 \end{pmatrix}\right), \tag{S37}$$

where, as described above, $F_M = F_0 + X_M + \epsilon_M - \epsilon_0$. Thus,

$$\mathbb{E}[X_M|F_0, F_M] = \frac{2M\sigma_a^2}{2M\sigma_a^2 + \sigma_\epsilon^2}\left(F_M - F_0\right). \tag{S38}$$

Thus, the contribution of additive effects to the conditional fitness gain linearly increases with $F_M - F_0$. For larger $F_M - F_0$, each additive effect is more likely to be positive.

### S4 Ascertained summary statistics in the LK fitness landscape

The LK model is somewhat impervious to analytic treatment under biased sampling of the global landscape. Thus, we treat only the statistics of a randomly ascertained landscape.

#### S4.1 Variance explained

For completeness, we provide a derivation of the amplitude spectrum, $A_m$, of the global landscape. We then solve for $A_m$ under down-sampling, where each of the $M$ focal loci are sampled uniformly from among the $L$ total loci. In solving for the down-sampled $A_m$, we obtained two distinct expressions, ultimately shown to be equivalent.

##### S4.1.1 Amplitude spectrum of the global landscape

We first derive the amplitude spectrum, $A_m$, of the global landscape. To begin, we consider the squared effect of an interaction term $\beta_{\mathbf{m}}$, where $\mathbf{m} \in \{0,1\}^L$, $|\mathbf{m}| = m$ indicates the loci involved in the interaction,

$$\begin{aligned}
\mathbb{E}[\beta_{\mathbf{m}}^2] &= 2^{-2L} \sum_i \sum_j \mathbb{E}[f_\ell^i f_\ell^j x_{j\mathbf{m}} x_{i\mathbf{m}}] \\
&= 2^{-2L} 2^L 2^{L-K} \binom{L-|\mathbf{m}|}{K-|\mathbf{m}|} \binom{L}{K}^{-1} \\
&= 2^{-K} \binom{L-|\mathbf{m}|}{K-|\mathbf{m}|} \binom{L}{K}^{-1},
\end{aligned} \tag{S39}$$

where $f_\ell^i$ and $f_\ell^j$ are the contributions of neighborhood $\ell$ to the fitness of genotypes $i$ and $j$, respectively. The second line follows from considering the fact that for each set of loci defined by $\mathbf{m}$, only neighborhoods which contain those $m$ loci will yield non-zero contributions to $\beta_{\mathbf{m}}^2$ in expectation.

The amplitude spectrum, $A_m$, is found by multiplying Eq. S39 by the number of interactions of order $m$,

$$\begin{aligned}
\mathbb{E}[A_m] &= 2^{-K} \binom{L-m}{K-m} \binom{L}{K}^{-1} \binom{L}{m} \\
&= 2^{-K} \frac{(L-m)!}{(L-K)!(K-m)!} \frac{K!(L-K)!}{L!} \frac{L!}{(L-m)!m!} \\
&= 2^{-K} \frac{K!}{(K-m)!m!} = 2^{-K} \binom{K}{m},
\end{aligned} \tag{S40}$$

in agreement with Neidhart et al. (2013).

##### S4.1.2 Amplitude spectrum of sub-sampled landscape

We now find the amplitude spectrum of a combinatorial library constructed from a set of $M$ uniformly sampled mutations, $\mathcal{M}$. We first consider the expectation of a squared first order effect, $\beta_\ell^2$.

**Derivation 1.** For $\ell \in \mathcal{M}$, a neighborhood $i$ will only contribute to $\beta_\ell^2$ when $\ell \in \mathcal{B}_i$. Unlike in the global landscape, for a given $i$, the number of sequences which contribute to $\beta_\ell^2$ is now determined by the number of loci $j \in \mathcal{M}$, $j \neq \ell$, for which $j \in \mathcal{B}_i$. Let $B_i$ be the number of loci in $\mathcal{M}$ in the neighborhood $\mathcal{B}_i$. Thus, we have, for $M < K$,

$$\begin{aligned}\mathbb{E}[\beta_\ell^2] &= \frac{1}{2^{2M}}\frac{1}{L}\sum_{i=1}^{L}\sum_{\mathbf{x}}\sum_{\mathbf{x}'}\mathbb{E}[f_i(\mathbf{x})f_i(\mathbf{x}')x_\ell x'_\ell] \\ &= \frac{1}{2^{2M}}\frac{1}{L}\sum_{i=1}^{L}\mathbb{E}\left[\mathbb{E}\left[\sum_{\mathbf{x}}\sum_{\mathbf{x}'}f_i(\mathbf{x})f_i(\mathbf{x}')x_\ell x'_\ell|B_i = j\right]\right],\end{aligned} \tag{S41}$$

and thus,

$$\begin{aligned}\mathbb{E}[\beta_\ell^2] &= \frac{1}{2^{2M}}\cdot 2^M\sum_{j=1}^{M}2^{M-j}\binom{M-1}{j-1}\binom{L-M}{K-j}\binom{L}{K}^{-1}\sigma^2 \\ &= \sigma^2\sum_{j=1}^{M}2^{-j}\binom{M-1}{j-1}\binom{L-M}{K-j}\binom{L}{K}^{-1},\end{aligned} \tag{S42}$$

with,

$$A_1 = \sigma^2 M\sum_{j=1}^{M}2^{-j}\binom{M-1}{j-1}\binom{L-M}{K-j}\binom{L}{K}^{-1}. \tag{S43}$$

For $m > 1$, a neighborhood $i$ will contribute to $\beta_{\mathbf{m}}^2$ only when all elements of $\mathbf{m}$ are in $\mathcal{B}_i$. Thus, we have

$$A_m = \sigma^2\binom{M}{m}\sum_{j=m}^{M}2^{-j}\binom{M-m}{j-m}\binom{L-M}{K-j}\binom{L}{K}^{-1}, \tag{S44}$$

where the factor of $2^{-j}$ comes from the fact that for a neighborhood containing the $m$ focal loci and $j-m$ non-focal loci (but still loci in the set $\mathcal{M}$), for each of the $2^M$ sequences there are $2^{M-j}$ sequences with the same set of states at the $m$ focal loci.

**Derivation 2.** Similarly, we can think of the variance explained under random ascertainment as a projection of the amplitude spectrum of the global landscape onto an $M$-dimensional subspace. In the global landscape, $\mathbb{E}[A_\ell] = 2^{-K}\binom{K}{\ell}\sigma^2$ (Eq. S40). An $\ell$-way interaction appears as an $m$-way interaction in the library if exactly $m$ of the $\ell$ loci are inside the library. The probability of this event is,

$$\mathbb{P}\{m - \text{way local}|\ell - \text{way global}\} = \binom{M}{m}\binom{L-M}{\ell-m}\binom{L}{\ell}^{-1}. \tag{S45}$$

Using Eq. S45, we then express the sub-sampled library amplitude spectrum, $A_m^{(M)}$, as a function of the full amplitude spectrum, $A_\ell$,

$$\begin{aligned}\mathbb{E}[A_m] &= \sum_{\ell=m}^{K}\mathbb{E}[A_\ell]\binom{M}{m}\binom{L-M}{\ell-m}\binom{L}{\ell}^{-1} \\ &= 2^{-K}\sigma^2\binom{M}{m}\sum_{\ell=m}^{K}\binom{K}{\ell}\binom{L-M}{\ell-m}\binom{L}{\ell}^{-1}.\end{aligned} \tag{S46}$$

In summary, the $\ell$-way interactions in the global landscape are converted to $m$-way interactions in the local landscape with probability given by the hypergeometric distribution. We reconcile these two perspectives below.

##### S4.1.3 Equivalence of the two expressions for the amplitude spectrum of a random LK library

We prove that the two expressions for the amplitude spectrum of a random library of size $2^M$ given in Eq. S44 and S46 are equivalent. For ease of exposition, we restate each of these expressions here, denoting them $A_m^M$ and $A_m^C$, respectively:

$$A_m^M = \binom{M}{m} \sum_{j=m}^{\min(M,K)} 2^{-j} \binom{L-M}{K-j} \binom{M-m}{j-m} \binom{L}{K}^{-1} \tag{S47}$$

$$A_m^C = 2^{-K} \binom{M}{m} \sum_{\ell=m}^{K} \binom{L-M}{\ell-m} \binom{K}{\ell} \binom{L}{\ell}^{-1} \tag{S48}$$

We begin with Eq. S48,

$$\begin{aligned} A_m^C &= 2^{-K} \binom{M}{m} \sum_{\ell=m}^{K} \binom{L-M}{\ell-m} \binom{K}{\ell} \binom{L}{\ell}^{-1} \\ &= 2^{-K} \binom{M}{m} \sum_{\ell=m}^{K} \binom{L-M}{\ell-m} \frac{K!}{L!} \frac{(L-\ell)!}{(K-\ell)!} \frac{(L-K)!}{(L-K)!} \\ &= 2^{-K} \binom{M}{m} \sum_{\ell=m}^{K} \binom{L-M}{\ell-m} \binom{L-\ell}{K-\ell} \binom{L}{K}^{-1} \end{aligned} \tag{S49}$$

Next, we make the following substitutions: $a = L - M$, $b = M - m$, $n = K - m$, $r = \ell - m$, and $q = j - m$. Taking out a factor of $\binom{M}{m}\binom{L}{K}^{-1}$, Eq. S47 and S49 become,

$$A_m^M = \sum_{q=0}^{\min(b,n)} 2^{-(q+m)} \binom{a}{n-q} \binom{b}{q},$$

and,

$$A_m^C = 2^{-K} \sum_{r=0}^{n} \binom{a}{r} \binom{a+b-r}{n-r}. \tag{S50}$$

We then apply Vandermonde's identity $\binom{x+y}{z} = \sum_{p=0}^{z} \binom{x}{p}\binom{y}{z-p}$ to the rightmost binomial coefficient in Eq. S50,

$$A_m^C = 2^{-K} \sum_{r=0}^{n} \binom{a}{r} \sum_{p=0}^{n-r} \binom{b}{p} \binom{a-r}{n-r-p} = 2^{-K} \sum_{p=0}^{n} \binom{b}{p} \sum_{r=0}^{n-p} \binom{a}{r} \binom{a-r}{n-r-p} \tag{S51}$$

where switching the order of the summations alters the limits. We now use the fact that $\binom{a}{r}\binom{a-r}{n-r-p} = \binom{a}{n-p}\binom{n-p}{r}$ and the Binomial Theorem, i.e., $\sum_{r=0}^{n-p} \binom{n-p}{r} = 2^{n-p}$, to further simplify the above expression,

$$\begin{aligned} A_m^C &= 2^{-K} \sum_{p=0}^{n} \binom{b}{p} \binom{a}{n-p} 2^{n-p} \\ &= \sum_{p=0}^{n} 2^{-K+(K-m)-p} \binom{b}{p} \binom{a}{n-p} \\ A_m^C &= \sum_{p=0}^{\min(b,n)} 2^{-(m+p)} \binom{b}{p} \binom{a}{n-p} = A_m^M, \end{aligned}$$

where the change in the limit is due to the fact that $\binom{b}{p} = 0$ if $p > b$, completing the proof.

### S5 Simulating from random fitness landscape models

#### S5.1 House of Cards

In each trajectory simulation, the ancestral sequence is sampled uniformly on the $L$-dimensional hypercube, $\mathbf{x}_0 \in \{-1, 1\}^L$. To generate the distribution of fitness effects (*dfe*), the fitness values of all single mutants of $\mathbf{x}_0$ are sampled independently and identically distributed (*iid*) from a standard normal distribution. The next sequence in the adaptive trajectory is sampled uniformly from among the beneficial mutants, i.e., those for which $F(\mathbf{x}') - F(\mathbf{x}_0) \geq 0$. This procedure, generating the *dfe* and sampling a beneficial mutant, is repeated for $M$ generations. Due to finite genome size, a subset of trajectories will get stuck before taking $M$ steps. For a given landscape, an adaptive trajectory is attempted from the same ancestral sequence up to twenty times. If unsuccessful the replicate is discarded, with the consequence that the average ancestral fitness across simulations is slightly biased toward negative values.

To allow for comparisons among scenarios, the fitness landscape in a given beneficial or deleterious simulation is inherited from the corresponding trajectory simulation. Thus, the $M$ beneficial and deleterious mutants are sampled from the previously generated *dfe* of the ancestral sequence. The fitness values of all additional sequences in the combinatorial library are either inherited from the trajectory fitness landscape (if already observed) or sampled *iid* from a standard normal.

Likewise, the random landscape in a given replicate is generated, in part, from the fitness landscape already explored by the three ascertainment scenarios. Fitness values of unobserved sequences are sampled *iid* from a standard normal distribution.

Simulation of the conditional scenario, however, is made more efficient by reconstitution of the fitness landscape. In particular, we retain the ancestral sequence and associated fitness value, while assigning the fitness of the adapted sequence endpoint to a random sequence at distance $M$ from the ancestor. The fitness values of all $2^M - 2$ intermediate sequences in the library are then sampled *iid* from a standard normal distribution. We conduct $N = 100$ simulations for $M = 2, 4, 6$ with $L = 100$.

#### S5.2 Rough Mount Fuji

To initialize a Rough Mount Fuji (RMF) landscape, the $L$ additive effects $\lambda_\ell$ are sampled from a standard normal distribution. As in the HoC, an ancestral sequence is sampled uniformly on the $L$-dimensional hypercube. The effect of a mutation at locus $\ell$ in the ancestral sequence $\mathbf{x}_0$ is given by,

$$F(\mathbf{x}_\ell) - F(\mathbf{x}_0) = -2\lambda_\ell x_{0\ell} + \epsilon(\mathbf{x}_\ell), \tag{S52}$$

where $\epsilon(\cdot) \overset{iid}{\sim} \mathcal{N}(0, \sigma_\epsilon^2)$, with values simulated on the fly. The trajectory, beneficial, deleterious, and randomly ascertained sub-landscapes are otherwise generated as in the HoC.

Due to the distance-dependent correlations in the RMF, generating the conditional landscape requires a distinct approach. We conduct a brute force search of sequences at distances of $M$ mutations to identify a sequence with fitness sufficiently close to the adapted trajectory sequence. The fitness values, $F'$, of each of one thousand sequences in up to twenty random sampled batches is sequentially assessed for proximity to the target fitness, i.e., whether $|F' - F_M| \leq 10^{-3}\sqrt{L\sigma_a^2 + \sigma_\epsilon^2}$. If a sequence is not identified at the end of this procedure, the entire replicate is discarded.

## S5.3 LK

Each neighborhood $\mathcal{B}_i$, $i = 1, \ldots, L$ (Eq. 14) is generated by sampling $K$ loci uniformly without replacement for each of the $L$ neighborhoods. For computational efficiency, the fitness contributions, $f_i$, of each $K$-length subsequence are sampled on the fly, with $f_i(\cdot) \overset{iid}{\sim} \mathcal{N}(0, \sigma^2)$, with $\sigma^2 = 1$. The fitness of a given sequence is the normalized sum of all $L$ neighborhood contributions (Eq. 14). In a simulated replicate, all ascertainment scenarios share a single underlying landscape realization.

The *dfe* at $\mathbf{x}_0$, is given by

$$F(\mathbf{x}_\ell) - F(\mathbf{x}_0) = \frac{1}{\sqrt{L}} \sum_{\ell \in \mathcal{B}_i} f_i(\mathbf{x}_\ell) - f_i(\mathbf{x_0}), \tag{S53}$$

where $\mathcal{B}_i$, $|\mathcal{B}_i| = K$, is the $i^{\text{th}}$ neighborhood. All other scenarios are otherwise sampled as described above; in the conditional library, the fitness of the derived sequence is required to be within $\epsilon = 10^{-3}$ of the trajectory endpoint fitness.

We simulate $N = 1{,}000$ simulations with $M = 4$, $L = 100$, with $1 \leq K \leq 100$.

### S6 Ascertained summary statistics in empirical landscapes

We provide additional methodological detail and support for claims made in the main text in the analysis of three empirical combinatorial mutagenesis experiments: (1) GB1 binding IgG-Fc (Wu et al., 2016); and ParD3 binding (2) cognate, ParE3, and (3) non-cognate, ParE2, toxins (Lite et al., 2020).

#### S6.1 Simulating ascertainment

In the random scenario, a mutation at each site was sampled uniformly from among the twenty possible states. In the beneficial and deleterious scenarios, the most and least fit single mutants at each site were selected, respectively. At each step in the adaptive trajectory, a mutation was sampled from the set of beneficial mutations at sites which had not already been mutated. In the conditional scenario, a mutant was chosen from the set of mutants for which (1) the phenotypic distance was within $\epsilon = .01$ of $F_M$, the endpoint fitness in the adaptive trajectory, and (2) the genetic distance from the ancestral sequence was equal to $M$.

#### S6.2 Data processing

For GB1, fitness values were estimated as the log-transformed ratio of post-selection, $N_1$, and initial, $N_0$, read counts,

$$F(\mathbf{x}) = \log\left(\frac{N_1(\mathbf{x}) + \eta}{N_0(\mathbf{x}) + \eta}\right), \tag{S54}$$

where $\eta = 1$ is a pseudo-count. For the ParD3 datasets, fitness values were obtained from Park et al. (2024) (see Lite et al. (2020) for more details).

#### S6.3 Estimating a global regression model

For each CME, the entire library was used to fit regression models of increasing order $m = 1, \ldots, M$ with all fitness estimates scaled to variance one (but not centered). Variance explained by the model was also evaluated with respect to the entire dataset.

The effect of substituting allele $a$ at locus $\ell$, $\hat{\beta}_{\ell(a)}$, was estimated independently for each locus-allele combination as,

$$\hat{\beta}_{\ell(a)} \equiv \langle F(\mathbf{x}) \rangle_{x_\ell = a} - \bar{F}, \tag{S55}$$

where $\langle F(\mathbf{x}) \rangle_{x_\ell = a}$ denotes an average over those genotypes with allele $a$ at locus $\ell$, and $\bar{F}$ is the average fitness of the library. The predicted phenotype of genotype $\mathbf{x}$ is then,

$$\hat{F}(\mathbf{x}) = \bar{F} + \sum_{\ell=1}^{M} \sum_{a \in \mathcal{A}} \hat{\beta}_{\ell(a)} \mathbb{1}\{x_\ell = a\}, \tag{S56}$$

where $\mathcal{A}$ is the set of all possible alleles.

#### S6.4 Robustness to measurement noise

In the main text, we suggested that the differences in the $R^2$ statistics between the trajectory, beneficial, and conditional scenarios with the random and deleterious scenarios could be primarily attributed to measurement noise. The rationale is that sampling deleterious mutations is likely to enrich for low fitness mutants, with fitness at or below the measurement detection threshold. At this lower limit, fitness becomes inversely proportional to the number of initial reads, which, if uncorrelated with fitness, implies that the fitness differences between such mutants are merely due to noise. Therefore, the $R^2$ statistics of the deleterious libraries will approximate that of the House of Cards (HoC) model, which assumes that the fitness values of each mutant are *iid* normal. As most mutants are deleterious, with a large mode at the detection threshold, the random library will also enrich, albeit less substantially, for low fitness mutants, akin to the deleterious library.

To test this hypothesis, we permuted the genotypes of the members of a given library and re-estimated the amount of variance explained by interactions of each order $m = 1, \dots, M-1$, $\tilde{A}_m$, $B = 100$ times. For each scenario, we then asked whether the observed $\tilde{A}_m$ statistic differed from that of the permuted sample by computing a two-sided empirical $p$-value,

$$p_m \equiv 2 \times \min \left\{ \frac{1 + \sum_{b=1}^{B} \mathbb{1}\{\tilde{A}_m > \tilde{A}_{mb}\}}{B+1}, \frac{1 + \sum_{b=1}^{B} \mathbb{1}\{\tilde{A}_m < \tilde{A}_{mb}\}}{B+1} \right\}, \tag{S57}$$

where $\tilde{A}_{mb}$ is the $\tilde{A}_m$ statistic of the $b^{\text{th}}$ permuted replicate. As suspected, all but the deleterious scenario deviated from the permuted distributions for $\tilde{A}_m$, $m = 1, \dots, M-1$ (Fig. S7).

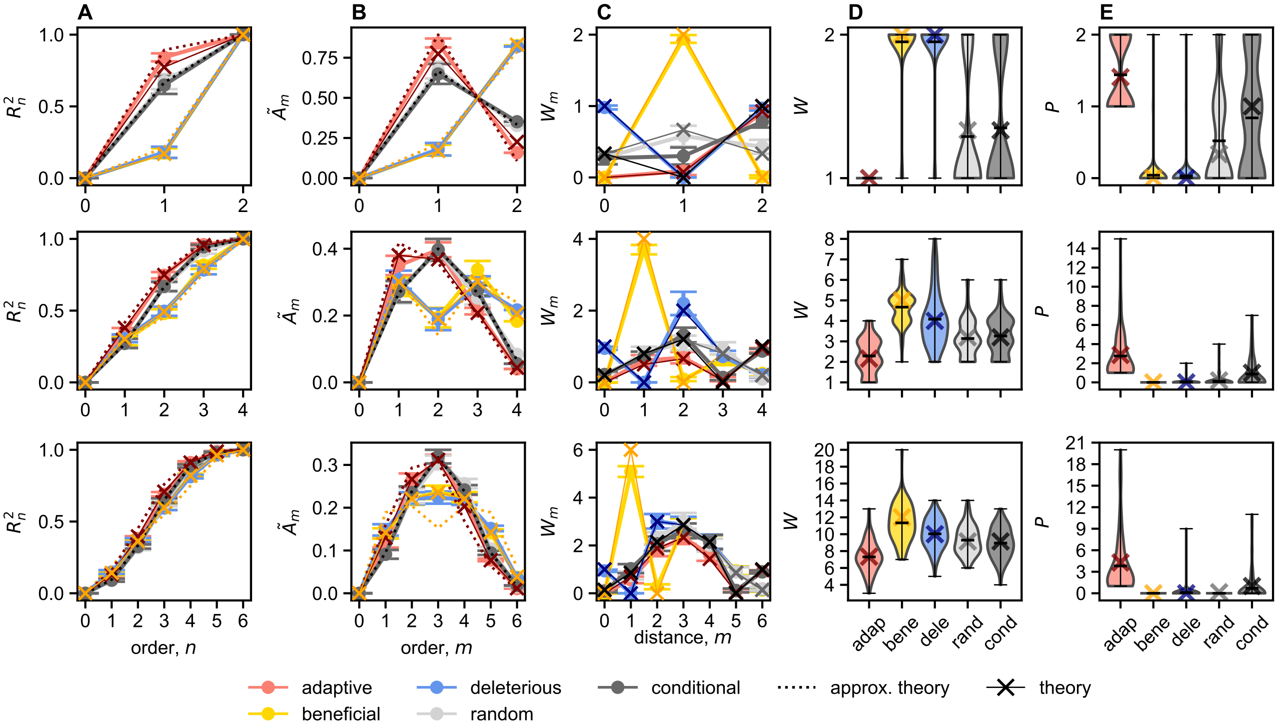

**Fig. S1. Statistics of ascertained libraries in House of Cards (HoC) landscapes.** **(A)** Variance explained by models of order $n$, $R_n^2$, $n = 0, 1, \ldots, M$, for combinatorial libraries of size $2^M$ ($M = 2, 4, 6$, top, middle, and bottom rows, respectively) for the adaptive trajectory (salmon), beneficial (yellow), deleterious (blue), random (gray), and conditional (dark gray) ascertainment schemes in HoC landscapes with $L = 100$. Each point is an average of $N = 100$ simulations. Error bars represent 95% confidence intervals. We also show theoretical expectations for $R_n^2$ in each scenario. The dotted line denotes expectations for $R_n^2$ in the trajectory, beneficial, and deleterious scenarios derived under the assumption that the atypical mutants in these libraries are *iid* samples from a distribution with mean $\mu_g = 4$ and $\sigma_g^2 = 4$, and deleterious single mutants from a distribution with mean $-\mu_g$ and variance $\sigma_g^2$. **(B)** The normalized amplitude spectrum, $\tilde{A}_m \equiv R_m^2 - R_{m-1}^2$ for each scenario, with all otherwise as in **A**. **(C)** The average number of local maxima as a function of distance, $m$, from the ancestral sequence, $W_m$. **(D)** Violin plots of the number of local maxima, $W$, and **(E)** accessible paths, $P$, across ascertainment schemes. We also show theoretical expectations for these quantities, as in **A**-**C**. Black hashes in the violin plots denote the maximum, mean, and minimum, respectively.

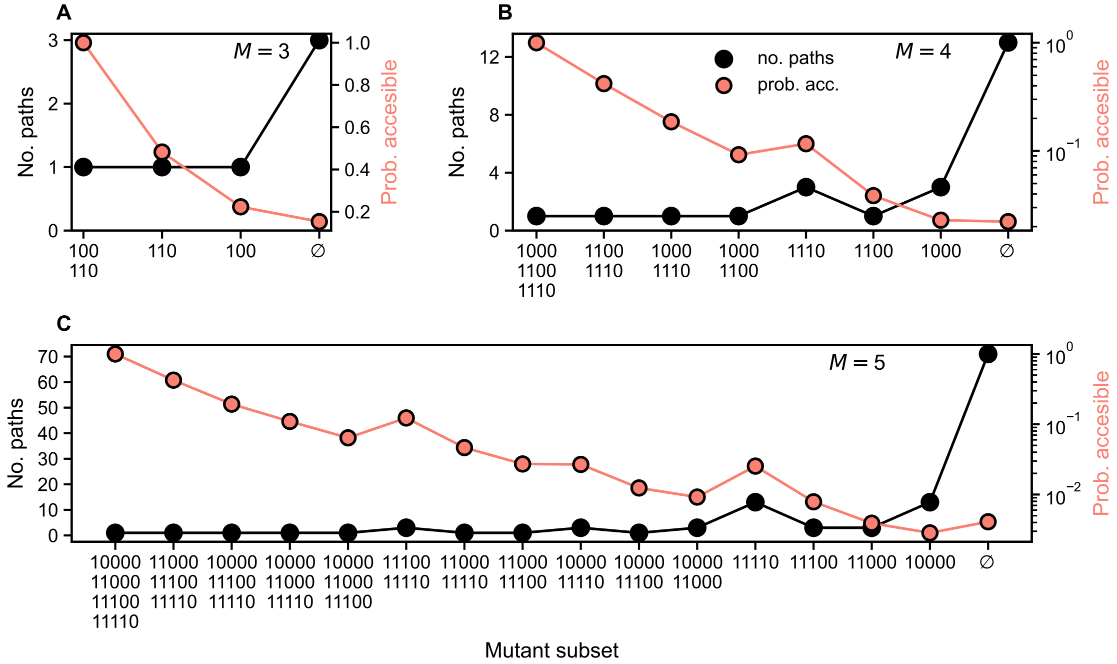

**Fig. S2. Number of paths which include mutant subsets.** The number of paths which include each subset of trajectory mutants (black points) for libraries constructed with **(A)** $M = 3$, **(B)** $M = 4$, and **(C)** $M = 5$ mutations which occurred along an uphill adaptive walk in a House of Cards (HoC) fitness landscape (y-axis, left). The empty set, $\emptyset$, refers to paths which contain no trajectory mutants outside of the ancestral and fully derived sequences. The salmon points denote the probability that a path containing a given subset is accessible (y-axis, right). The probabilities are computed using the average fitness values of mutants along adaptive trajectories of length $M$ ($N = 100$ simulations, $L = 100$).

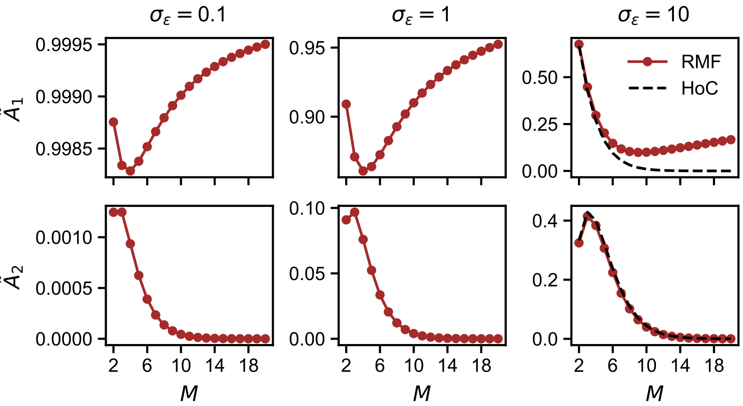

**Fig. S3. Amplitude spectra under random ascertainment in Rough Mount Fuji landscapes.** Analytic expectations for the variance explained by interactions of first and second order, $\tilde{A}_1$ (top row) and $\tilde{A}_2$ (bottom row), respectively, for increasing House of Cards (HoC) variance, $\sigma_\epsilon = .1, 1, 10$ (from left to right).

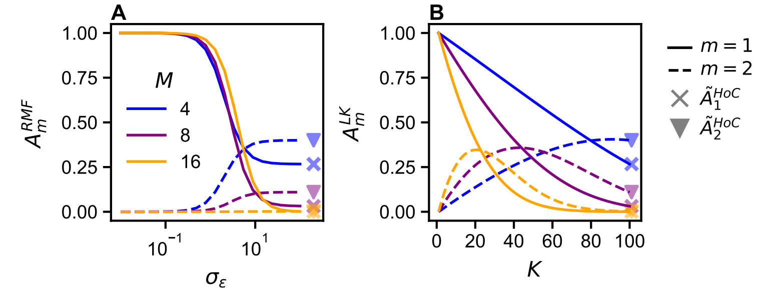

**Fig. S4. Amplitude spectra of random ascertainment in two random landscape models.** Analytic expressions for the normalized amplitude spectra, $\tilde{A}_m$, for $m = 1$ (solid lines) and $m = 2$ (dashed lines) for the **(A)** Rough Mount Fuji (RMF) and **(B)** LK landscapes for random libraries consisting of $M = 4, 8, 16$ mutations (colors) sub-sampled from global landscapes with $L = 100$. The House of Cards (HoC) expectations for $\tilde{A}_1$ and $\tilde{A}_2$ are denoted by the $\times$'s and triangles, respectively.

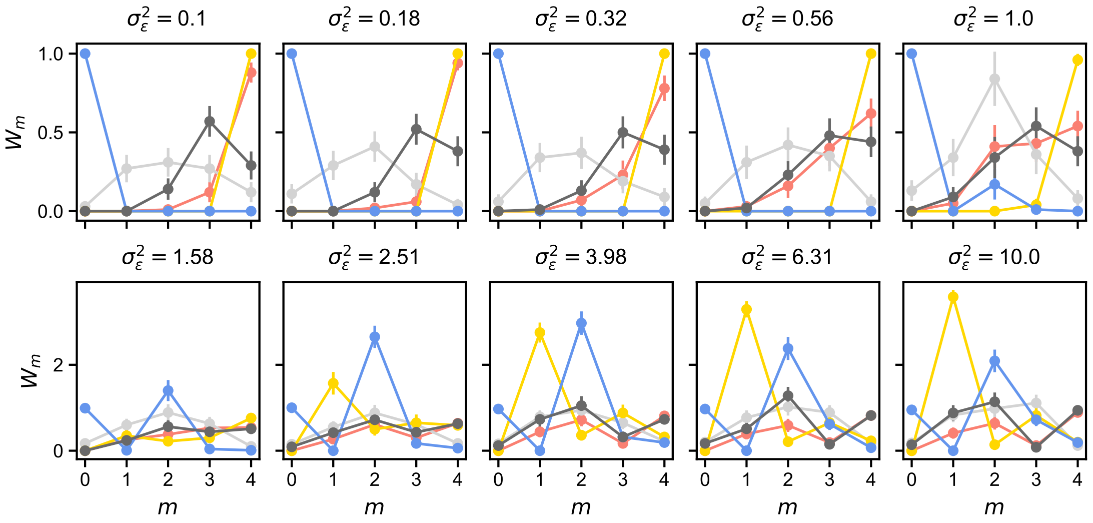

**Fig. S5. Number of local maxima as a function of distance from the ancestral sequence in Rough Mount Fuji landscapes.** Each subplot shows the average number of local maxima as a function of distance from the ancestor, $W_m$, for $m = 1, \ldots, 4$ in ascertained libraries of size $2^M$ with $M = 4$. The variance of the House of Cards (HoC) component, $\sigma_\epsilon^2$, increases from left to right by row. Each ascertainment scheme is indicated by different color. In all plots averages are computed from $N = 100$ simulations.

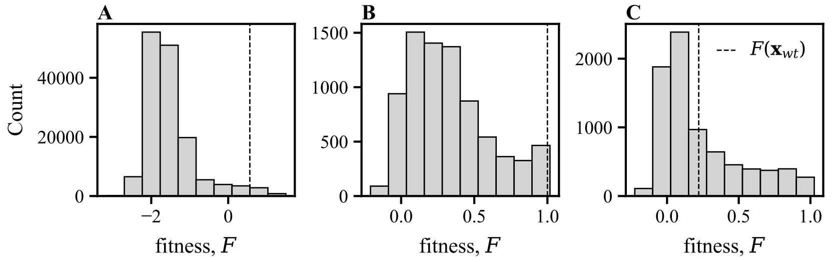

**Fig. S6. Distributions of observed fitness values in empirical mutagenesis experiments. (A)-(C)** Histograms of log or log relative fitness, $F$, in GB1 (Wu et al., 2016), ParD3 binding ParE3, and ParD3 binding ParE2 (Lite et al., 2020). The wildtype fitness in each case is indicated by the dashed line.

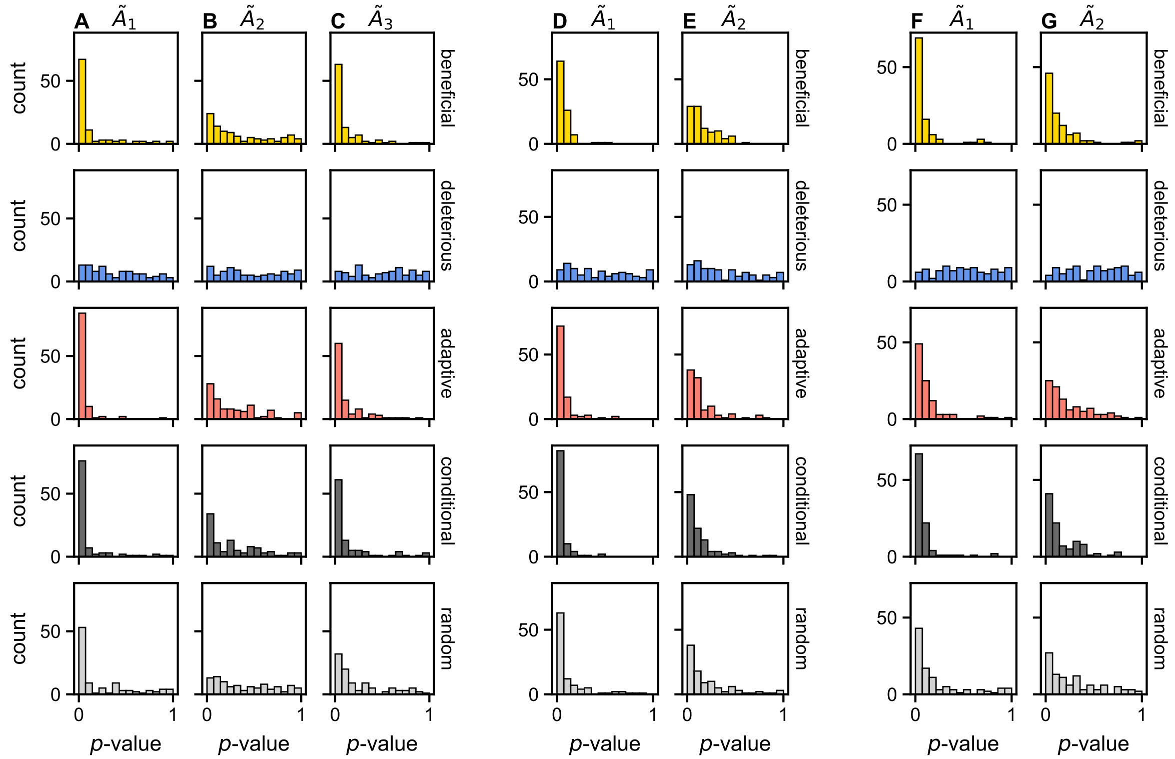

**Fig. S7. Permuted $R^2$ statistics.** The genotype labels in each ascertained library (rows) were permuted before recomputing the variance explained by interactions of a given order, $\tilde{A}_m := R_m^2 - R_{m-1}^2$. Each column displays histograms of the empirical two-sided $p$-values comparing the observed $\tilde{A}_m$ to the distribution of permuted statistics, for **(A-C)** GB1, $m = 1, 2, 3$, respectively; **(D-E)** ParD3-ParE3, $m = 1, 2$, respectively; **(F-G)** ParD3-ParE2, $m = 1, 2$, respectively.

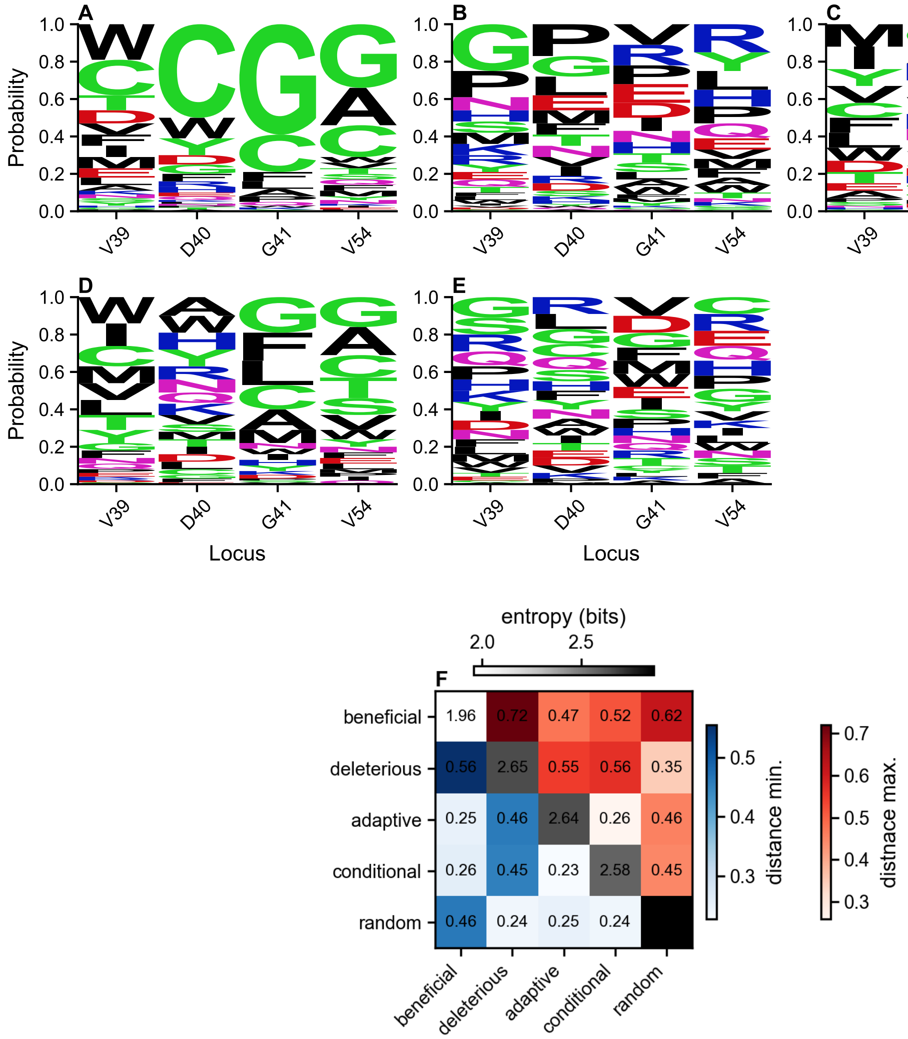

**Fig. S8. Sequence logos for each ascertainment scenario in the GB1 dataset.** Sequence logos representing amino acid occurrence at each of the $M = 4$ positions mutated by Wu et al. (2016) across $N = 100$ replicates and ascertainment scenarios: **(A)** beneficial, **(B)** deleterious, **(C)** adaptive trajectory, **(D)** conditional, and **(E)** random. The height of each amino acid letter corresponds to the frequency at which that amino acid is found at site $m$ in a given ascertainment scenario. **(F)** The minimum (lower triangle, blue) and maximum (upper triangle, gray) Jensen-Shannon distance between the sequence logos of each pair of ascertainment scenarios. The Shannon entropy of the sequence logo of each scenario is given on the diagonal in gray.

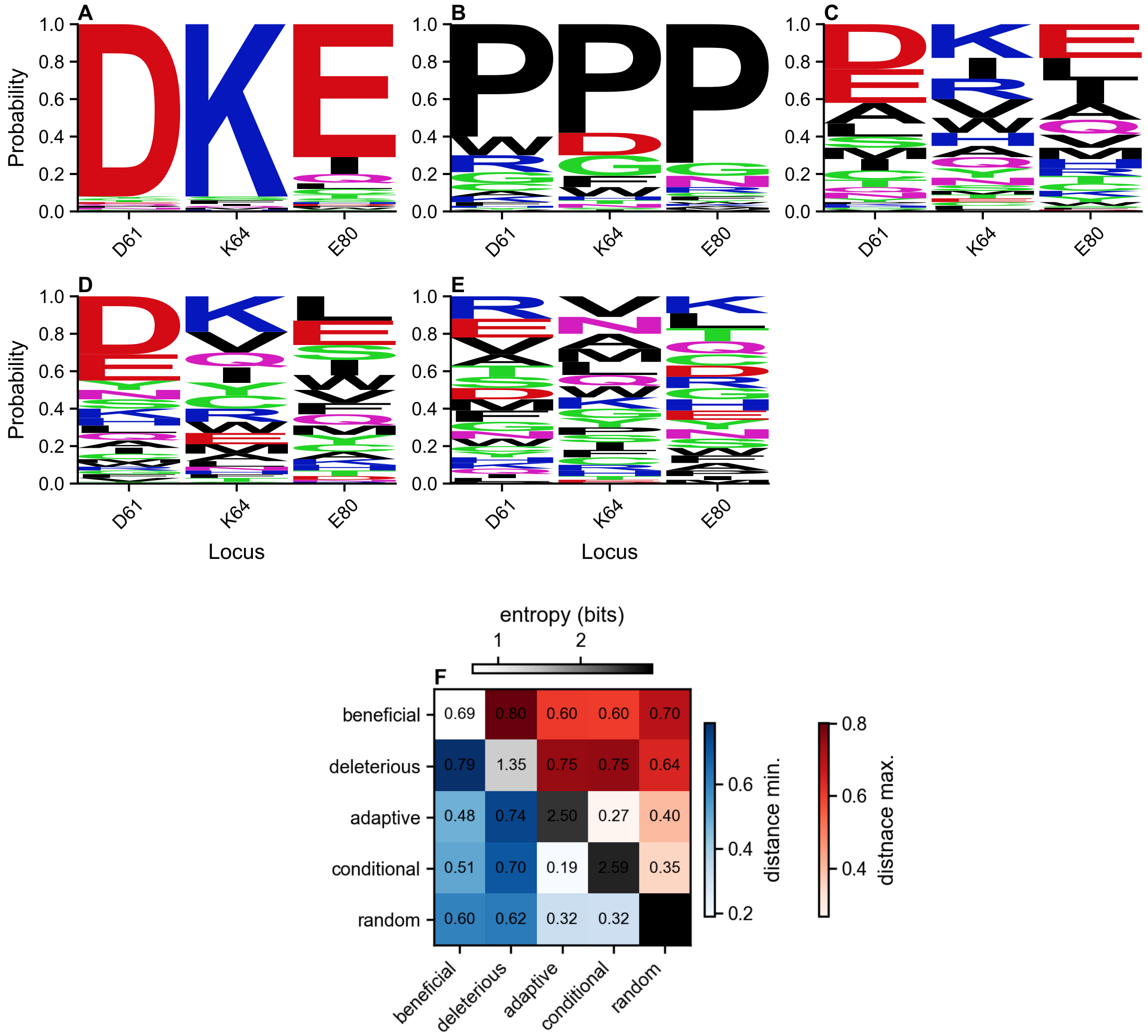

**Fig. S9. Sequence logos for each ascertainment scenario in the ParD3-ParE3 dataset.** All is as in Fig. S8, except here for sub-sampling of the ParD3-ParE3 combinatorial mutagenesis dataset (Lite et al., 2020).

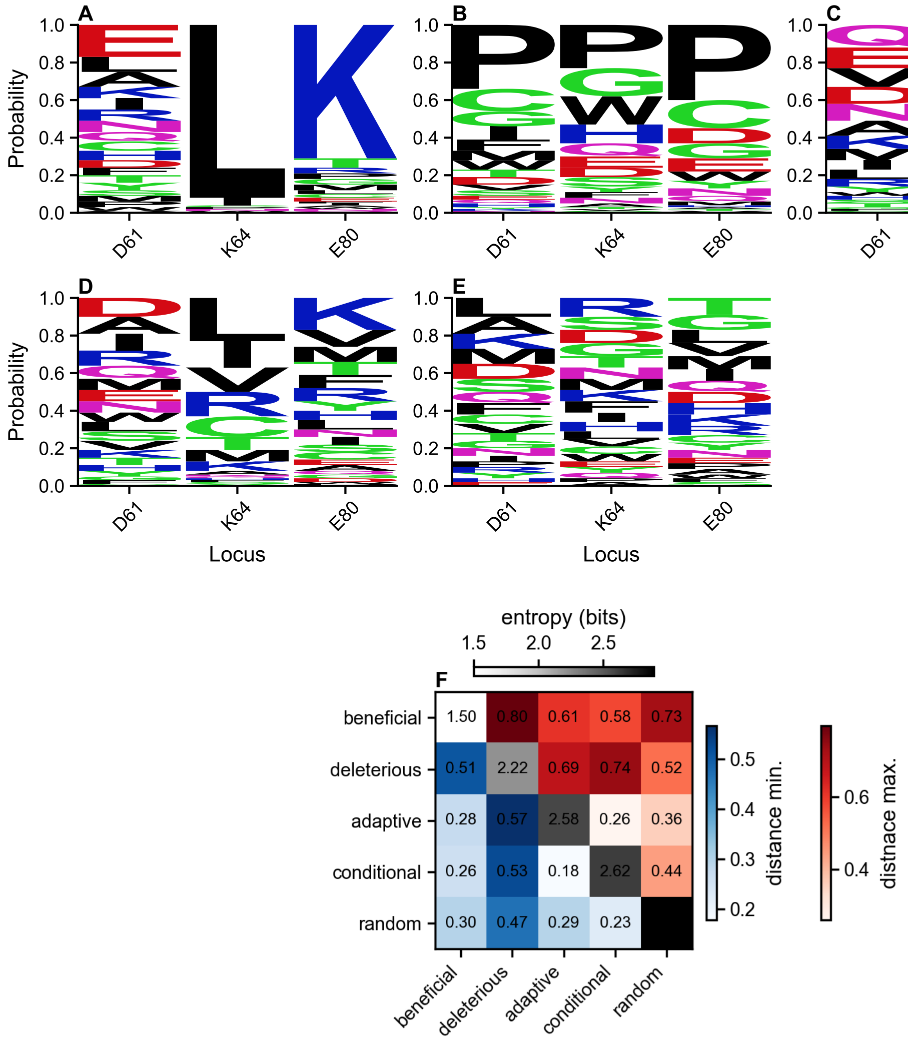

**Fig. S10. Sequence logos for each ascertainment scenario in the ParD3-ParE2 dataset.** All is as in Fig. S8, except here for sub-sampling of the ParD3-ParE2 combinatorial mutagenesis dataset (Lite et al., 2020).

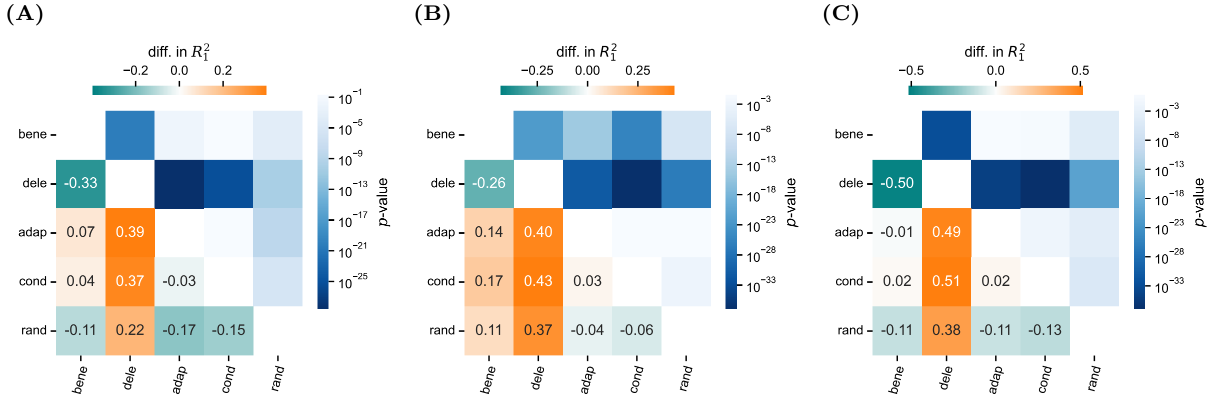

**Fig. S11. Differences in $R_1^2$ among scenarios in each of the three datasets.** In **(A)**-**(C)**, the lower triangle of each heatmap shows the average difference in $R_1^2$ between the scenarios and the upper triangle the corresponding $p$-value. **(A)** GB1 (Wu et al., 2016); **(B)** ParD3-ParE3; and **(C)** ParD3-ParE2 (Lite et al., 2020).

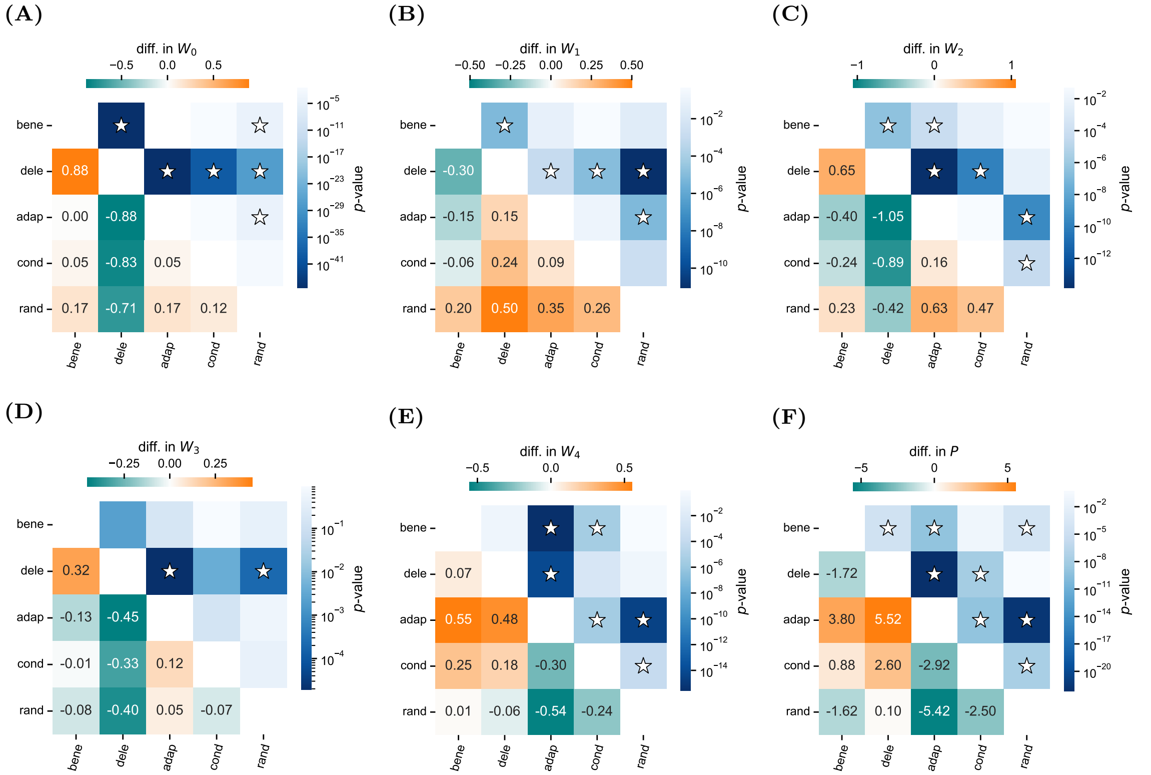

**Fig. S12. Differences in the incidence of local maxima and number of accessible paths among scenarios in GB1.** In **(A)**-**(E)**, the lower triangle of each heatmap shows the average difference in the number of local maxima at distance $m$ from the ancestral sequence, $W_m$ ($m = 0, \ldots, M$), between the scenarios and the upper triangle the corresponding $p$-value. Stars indicate statistical significance after a Bonferroni multiple test correction applied across all observed $W_m$ statistics ($\alpha = 0.05/(10 \times (M+1))$). **(F)** The same as in **(A)**-**(E)** except for the number of accessible paths, $P$ ($\alpha = 0.05/10$).

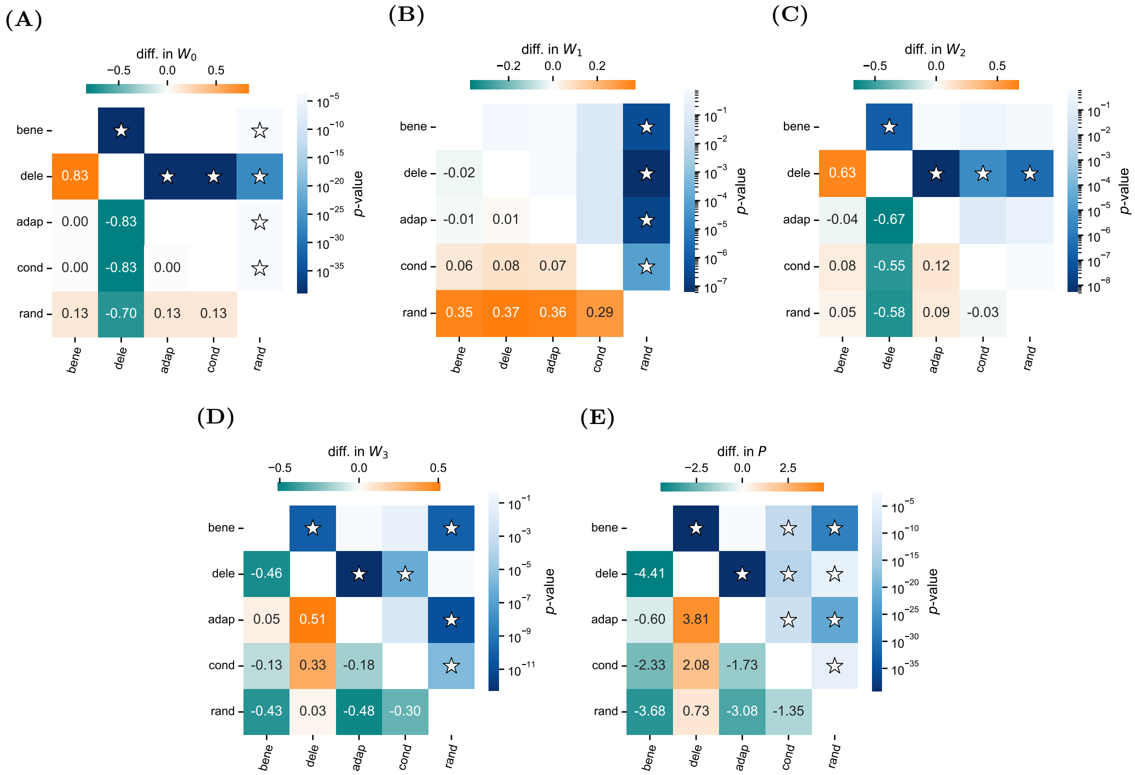

**Fig. S13. Differences in the incidence of local maxima and number of accessible paths among scenarios in ParE3.** The same as in Fig. S12 except for ParD3-ParE3 and with **(A)**-**(D)** corresponding to $W_m$, $m = 0, \ldots, 3$, and $P$ in **(E)**.

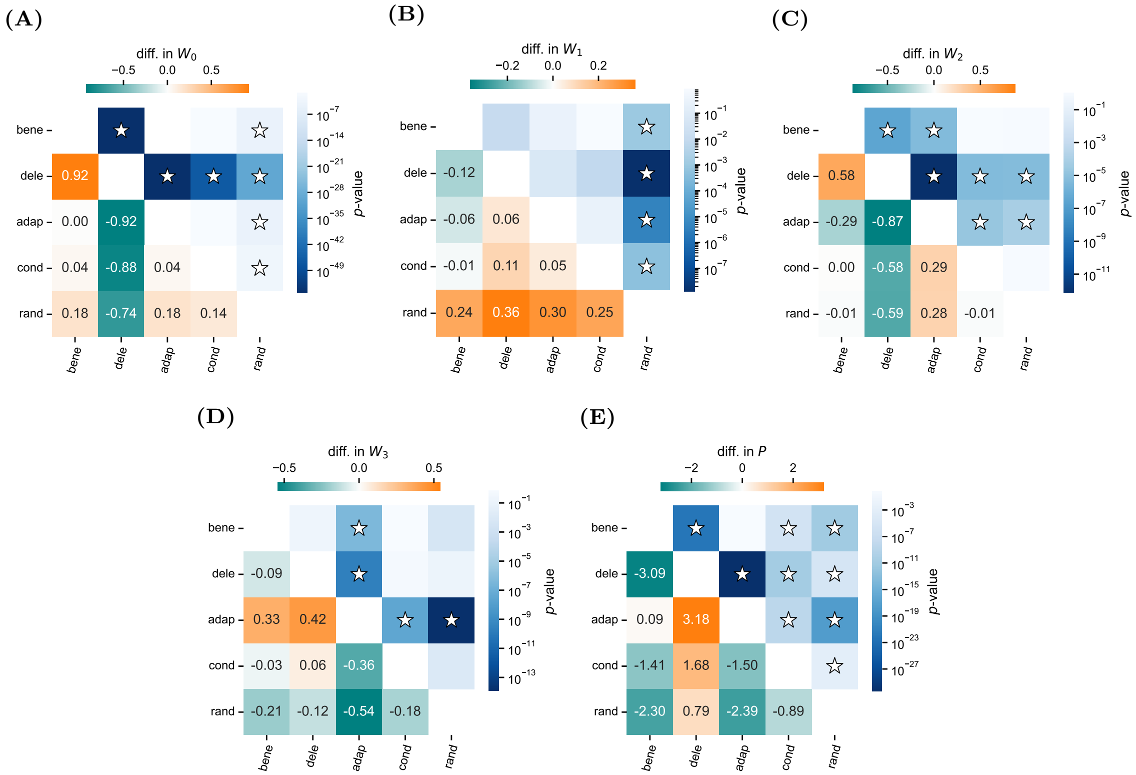

**Fig. S14. Differences in the incidence of local maxima and number of accessible paths among scenarios in ParE2.** The same as in Fig. S13 except for ParD3-ParE2.
